# Chromosome-scale genome assembly and gene annotation of the hydrothermal vent annelid *Alvinella pompejana* yield insight into animal evolution in extreme environments

**DOI:** 10.1101/2024.06.25.600561

**Authors:** Sami El Hilali, Philippe Dru, Alan Le Moan, Yang I Li, Martijn A. Huynen, André Hoelz, Robert C. Robinson, José M. Martín-Durán, Didier Jollivet, Adam Claridge-Chang, Richard R. Copley

## Abstract

The types of genomic change needed for environmental adaptation are of great interest. Annelid worms are a large phylum found in a rich diversity of habitats, giving opportunities to explore this issue. We report the chromosome level genome sequence of the Pompeii worm, the annelid *Alvinella pompejana*, an inhabitant of an extreme deep-sea hydrothermal vent environment. We find strong but heterogeneously distributed genetic divergence between populations taken from either side of the equator. Using transcript data, we produced a set of gene models and analysed the predicted protein set in the light of past hypotheses about the thermotolerance of *Alvinella*, and compared it to other recently sequenced annelid vent worms. We do not find evidence of a more extreme genome wide amino acid composition than other species, neither do we find evidence for rapid genome evolution in the form of disrupted synteny. We discount the hypothesis of loss of amino acid biosynthesis genes associated with obligate symbioses reported in siboglinid annelids. We do find evidence of a parallel increase in the number of globin encoding genes and loss of light sensitive opsins and cryptochromes. *Alvinella* encodes several respiratory enzymes unusual for bilaterian animals, suggesting an ability to better tolerate hypoxic environments.

## Introduction

The Pompeii worm, *Alvinella pompejana* Desbruyères & Laubier, inhabits the hottest part of the hydrothermal vent environment along the East Pacific Rise. At depths of several kilometres, these vents release superheated, metal rich water, potentially exposing *Alvinella* to toxic chemicals and extreme temperature and pressure. The thermal optimum for *A. pompejana* is likely to be beyond 42°C but below 50°C [1], and the related alvinellid *Paralvinella sulfincola* has been shown to prefer temperatures in the 40-50°C range and tolerate 55°C for brief periods [2]. These compare with body temperatures in excess of 45°C that have been reported in some species of birds [3,4] and a further caveat is that there is likely to be an interplay between behaviour and the fluctuating temperatures of vent environments [5].

In part owing to their inaccessibility, molecular work on alvinellids has been limited, but relatively large transcript sets have been constructed [6,7] and *Alvinella pompejana* has been proposed as a source of stable metazoan proteins suitable for structural and functional studies [7–13]. Attempts to correlate protein amino acid composition with thermotolerance have been a recurring theme e.g. [7,14]. Most recently, Jollivet and co-workers performed transcriptome sequencing of 6 alvinellid species demonstrating that the ancestral alvinellid was likely thermophilic and that cold-water adapted alvinellids showed distinct amino acid preferences [15].

Alvinellids are members of the terebellid clade of annelids [16]. Another kind of annelid, the vestimentiferan tubeworms from the siboglinid family are also found in deep sea hydrothermal vents and cold seeps, but live in colder habitats. Recently, the genomes of three of these siboglinids have been reported [17–19], and that of a fourth non-vestimentiferan, the idiosyncratic bone-eating *Osedax frankpressi* [20]. Siboglinids are unusual among animals in lacking a digestive system. Instead they have a trophosome, housing symbiotic bacteria that oxidize hydrogen sulfide to generate ATP and supply nutrients to the host. Alvinellids, in contrast, are grazers with an apparently intact digestive system [21] and epibiotic symbionts [22]. Terebellid and siboglinid annelids are not closely related [23], suggesting that their colonisation of deep sea vents has occurred independently, and raising the question of how their strategies to adapt to the deep sea vent environment differ.

Recently, a fragmented whole genome sequence of *Alvinella pompejana* was analysed together with its transcriptome to explore the genomic patterns in the diversification of *A. pompejana* and the closely related *A. caudata* [24]. A sub-chromosome level genome of *Paralvinella palmiformis* has been reported and analysed within the context of an epigenome study, without extensive study of its protein coding gene set [25]. Here we report a chromosome level genome assembly of *Alvinella pompejana*. We study the genome-wide population structure of resequenced individuals collected from two distinct genetic units on each side of the Equator [26,27] and note the existence of semi-permeable barriers to gene exchanges between these two units. We go on to compare the protein coding gene content of the reference genome to that from siboglinids and other relevant taxa. We show that its extreme habitat has led to both convergent and clade-specific changes, plausibly related to vent adaptation.

## Results

### Genome

Following extraction of high molecular weight genomic DNA, we sequenced and assembled a high quality genome from a single individual (see methods). Further scaffolding of the assembly using sequence from Hi-C and Chicago libraries (Dovetail Genomics, see methods) resulted in a 367.4 Mb genome (N50=21.45 Mb) (**Table S1**) out of which 95% (348.5 Mb) is covered by 17 pseudo-chromosomes (with scaffolds longer than 1 Mb) (**Table S2**). The size of the assembly is in line with previously reported measures using flow cytometry [28]. Dixon and co-workers report a karyotype of 16 chromosomes for *Alvinella pompejana,* although they note sampling difficulties and poor quality of chromosomes [29].

The genome shows a GC content of 43.3%, the highest level seen in annelids so far (**Table S1**). Its repetitive content is 23.3%, lower than other sequenced annelid genomes, with the exception of the ‘miniaturized’ genome of *Dimorphilus gyrociliatus* [30] (**Table S1**). Our assembly was 94.4% (BUSCO5 metazoa odb10) complete compared to the currently available metazoan gene set, in line with recently released high quality genomes of annelids. We used RNA-seq data to support gene predictions and produced a set of 19,179 gene models. After further curation of the genes (methods) we retained 18,739 protein coding genes.

### Mitochondrial genome

*Alvinella pompejana* tolerates a broad range of temperature [31]. A study conducted on marine invertebrates showed that the mitochondrion of deep-sea species living in zones with a high and highly variable temperatures was more resistant to high temperatures than that of other species living in cooler deep-sea and shallow water environments [32]. Mitochondrial respiration of *Alvinella* was reported to be the most stable when compared to other hot-adapted species, including the siboglinid *Riftia pachyptila* and the brachyuran crab *Bythograea thermydron*. We find the full mitochondrial genome of *Alvinella* consists of a 16.478 kb circular contig showing a GC content of 41%. Its relative order of protein coding and ribosomal RNA genes is identical to the recently released mitochondrial genomes of the siboglinid *Riftia pachyptila* [19], and the alvinellid *Paralvinella palmiformis* [33] and the highly conserved order in Pleistoannelida [34].

### Geographical differentiation of the genome

Genetic studies of Pompeii worm populations along the East Pacific Rise (EPR) showed two distinct genetic units on either side of the Equator [26,27], with the existence of a hybrid zone around 7°S/EPR. We confirmed this finding here using 3.33 million SNPs obtained after the WGS data processing. The genetic differentiation between individuals on either side of the Equator was much higher than expectation based on the divergence of genes [24] (**Figure 3a**), with an average 70% polymorphism identity between individuals of the same population but 50% between the southern and northern populations. The differentiation is, however, unevenly distributed with some portions of chromosome highly differentiated and others not (**Figure 3b**). Both individuals sampled in the south EPR are introgressed by alleles from the north, demonstrating that hybridization of the two geographic forms extends south of the hybrid zone described around 7°S/EPR. The introgression footprints are often located in different genomic regions across the two introgressed individuals (**Figure S1**). In addition, the introgressing alleles are distributed in blocks along the genome (**Figure 3b**). This pattern suggests the presence of non-recombining regions where long and divergent haplotypes are maintained for significant periods after their introgression. Chromosome 11 shows a remarkably contrasting pattern of divergence between the north and the south EPR (**Figure 3c**), with homogeneous divergence found over the entire chromosome, no signs of introgression, and lower genetic diversity (H_obs_ C11= 0.19 vs H_obs_ overall = 0.38). We suggest that this chromosome acts as major genetic barrier between the north and south EPR populations of *A. pompejana*.

**Figure 1.**
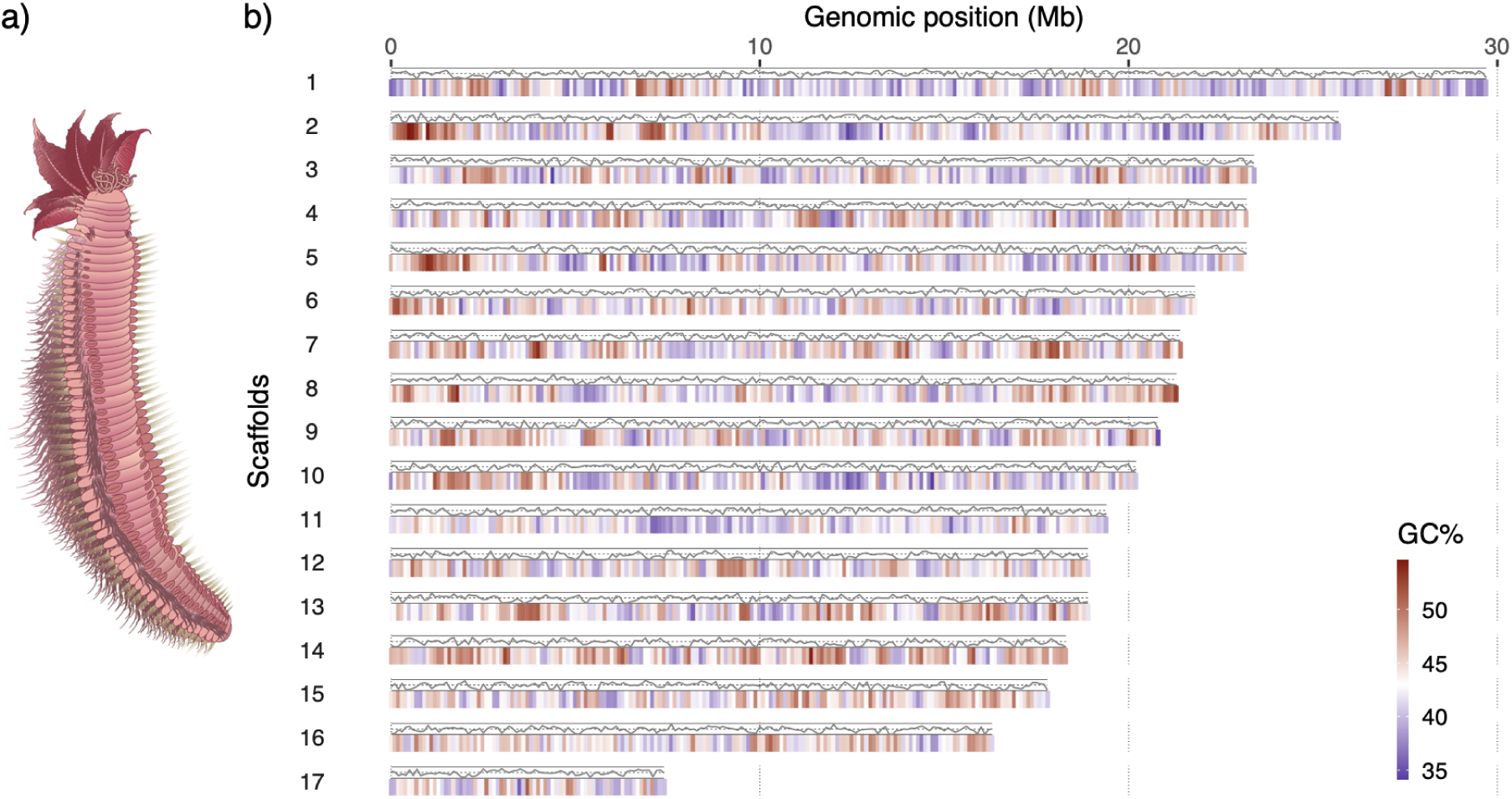
a) Schematic of *Alvinella pompejana* (∼10cm) b) Pseudo-chromosomes of *Alvinella* genome assembly. The GC content (red/blue colour ramp) and gene density (line) over 100kb windows are shown with the latter computed as the percentage of the coverage by predicted genes in 100kb non-overlapping windows, with the top line indicating 100%.

**Figure 2.**
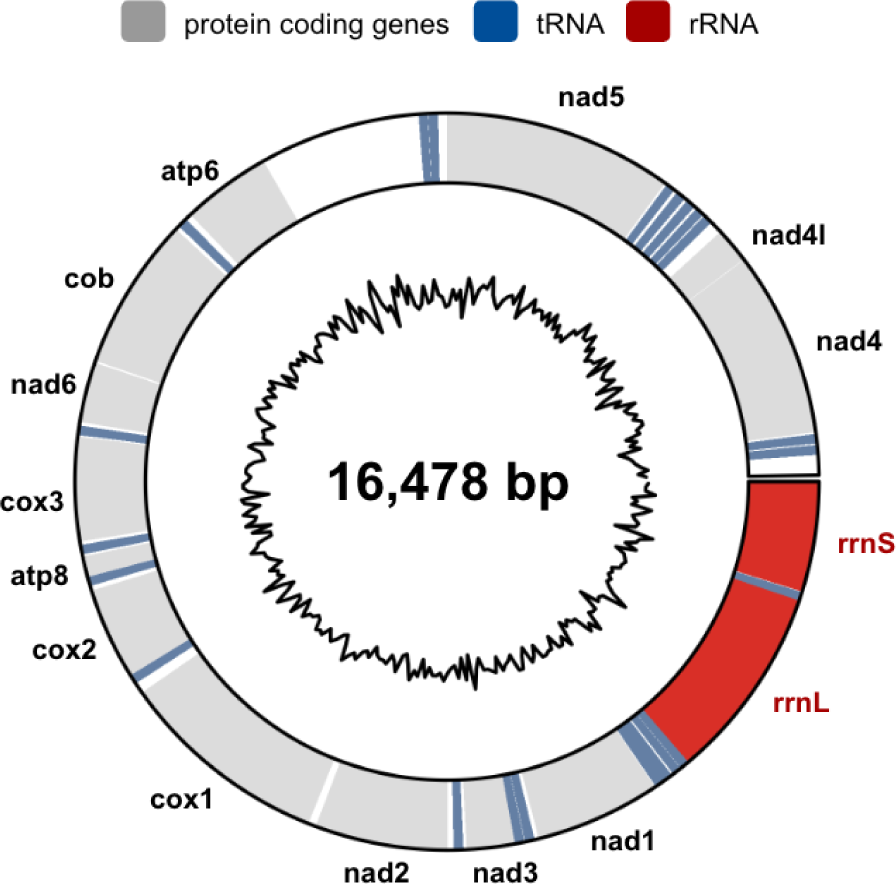
Fully assembled mitochondrial genome. The inner circle indicates the GC content calculated over adjacent genomic windows of 50 bp. Drawing using the circlize R package [35]

**Figure 3:**
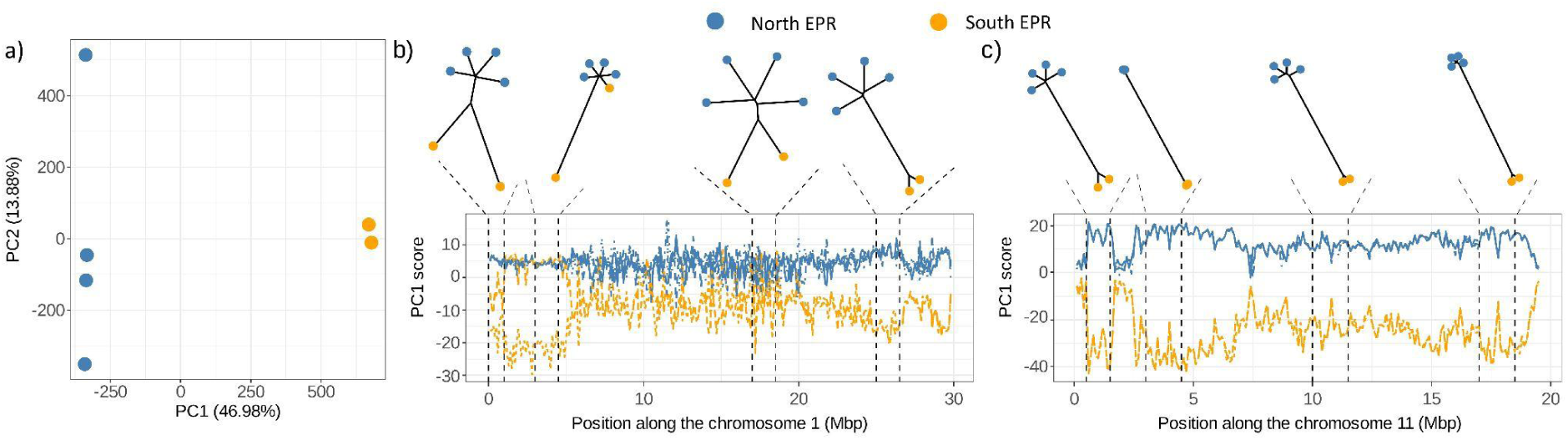
Genome-wide population structure of *A. pompejana* explored across two EPR South samples (in orange) and four EPR north samples (blue). The graph in a) shows a Principal Component Analysis (PCA) performed on the six samples genotyped at 3.33M SNPs. Most of the variation is captured by PC1 (PC1 = 46.98%; PC2 = 13.88%) and distinguishes samples from either side of the equator. Graphs in b) and c) show detailed analyses of genomics regions with contrasted patterns of divergence in b) chromosome 1 and c) chromosome 11. In each subplot, the top graphs show phylogenetic trees computed from the polymorphism data extracted from 1Mbp to 1.5Mbp windows defined from the patterns of divergence observed in the local PCA plot at the bottom. In the local PCA plots, each line follows the individual PC score of each sample over a sliding window. The vertical dotted lines show the limit of the genomics regions that were used to infer the phylogenetic trees. On chromosome 1, the phylogenies showed, in order of appearance, region of introgression where one EPR south samples is located at mid distance from the EPR north samples and the other EPR south sample, a region of introgression where one EPR south sample clustered perfectly with the EPR north samples, a region of weak divergence where all samples were similarly distant, and a region of high divergence where individuals on either side of the equator were clearly separated. On chromosome 11, all phylogenetic trees showed clear North – South EPR separation, with the second and last trees also showing small terminal branches expected from low genetic diversity within each population.

### Annelid phylogeny from complete genomes

We used phylobayes to calculate an annelid phylogeny using the CAT+GTR model of sequence evolution (**Figure 4**) [36]. The tree demonstrates the distinction between *Alvinella pompejana* and the siboglinids (*L. luymesi, P. echinospica, R. pachyptila*). In our analysis *Alvinella* is the sister of a *c*lade of the clitellate annelids (*H. robusta* and *E. andrei*) and *Capitella*. The branch lengths separating these clades are extremely short and have previously been resolved differently with different molecular datasets [37,38].

**Figure 4:**
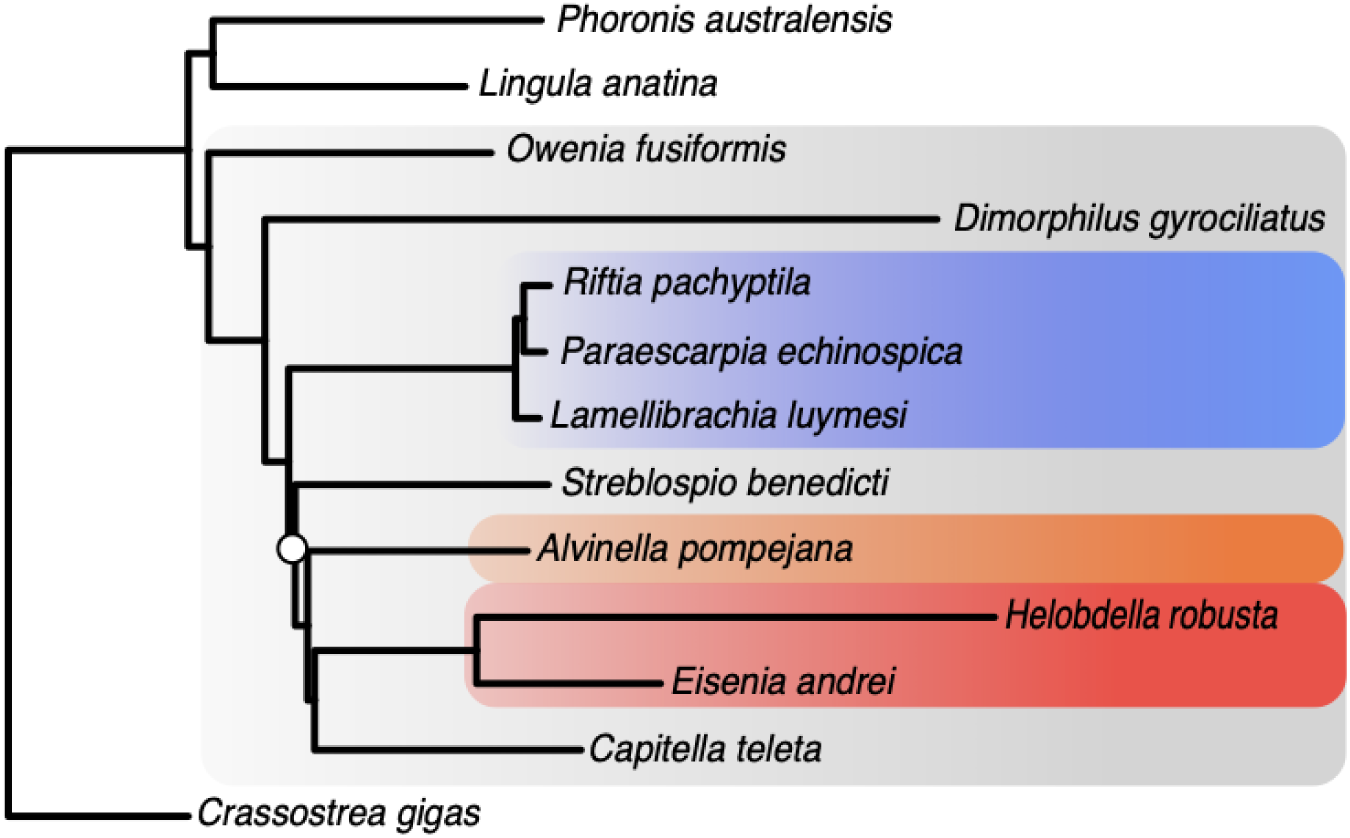
Phylobayes CAT+GTR phylogenetic tree of representative annelids with complete genomes. Calculated from a concatenated alignment of protein sequences corresponding to 301 orthologous genes from 10 annelid taxa and 3 outgroups (100,278 positions, see methods). The annelid clade has a grey background, and within that, siboglinids blue, clitellates red and *Alvinella* orange. *Lingula anatina* (brachiopod) and *Crassostrea gigas* (mollusc) are shown as outgroups. All nodes have a posterior probability of 1, except that of *S. benedicti* (0.75, marked with circle). Molluscs are placed as the root following [39].

The position of *Steblospio benedict*i was unstable, with 3 out of 4 chains showing the position in **Figure 4**, and the fourth, a position as sister to the clade of siboglinids, clitellates, *Alvinella* and *Capitella*. It is notable that neither the *Alvinella* nor siboglinid branch lengths show the rapid evolution of the clitellates or *Dimorphilus*.

### Evidence for a symbiont

Given the complexity of the vent ecosystem, and the prevalence of symbioses, we were mildly surprised to find that only 574 out of the original 19179 predicted protein sequences had a bacterial best hit when searched against the NR database of the NCBI (see methods). By far the majority of these (455 out of the 574) were encoded on sequences that did not form part of the 17 pseudo-chromosomes, although these non-pseudo-chromosomal sequences encode far fewer proteins (919 vs 18260 on the pseudo-chromosomes), so we suspect they represent bacterial contigs rather than genomically integrated horizontal gene transfer events. We sequenced a tissue sample taken from adult gills (see methods). As episymbionts are found mainly on the dorsal surface of the worm body, it seems likely that we have avoided bacterial sequencing to a large extent. In contrast, an earlier draft of the *A. pompejana* genome, assembled from different sequence reads, does appear to contain an extensive set of proteins from Bacteria (see methods and **Table S6**), with best hits including sequences labelled as *Alvinella pompejana* symbionts, epsilonproteobacteria (Campylobacterota), the sulfate reducing *Desulfobulbus* and metabolically significant genes such as dissimilatory sulfite reductase (**Table S7**) [24,40,41].

Halanych and co-workers presented an analysis of the presence and absence of amino acid biosynthetic genes in the siboglinid *Lamellibrachia luymesi* and concluded that it lacks many genes essential for amino acid biosynthesis, contrasting it with *Capitella teleta*, which they claimed encoded many of these missing genes [17]. This finding was repeated in the report of the genome sequence of *Riftia pachyptila* [19] and in both cases regarded as consistent with loss of the enzymes and a dependence of siboglinids on their bacterial symbionts for amino acids and co-factors. The absence of these amino acid biosynthetic pathways is, however, a synapomorphy of the Metazoa [42–45], so, if true, their reported presence in *Capitella* and hence inferred loss in the lineage leading to siboglinids would be a remarkably unparsimonious environmental adaptation.

We therefore reanalysed those putative *Capitella* protein sequences, without *Lamellibrachia* counterparts, that form the basis for the hypothesis. We found that most are very similar to bacterial proteins (**Table S4**), typically an *Endozoicomonas* species [46], but none have credible annelid, or in most cases, even animal orthologs (except in two cases where they appear to have been overlooked by Li *et al*.). Although it is possible that a proportion of these represent horizontal gene transfer into the *Capitella* genome (or an unknown symbiont), it seems likely that they are bacterial contaminants in the *Capitella* data [47]. If this interpretation is correct, it suggests that the repertoire of amino acid biosynthesis pathways in siboglinids is not exceptional among annelids or animals.

### Amino acid composition

The thermophily trait of the Pompeii worm has previously been proposed to rely on the amino acid composition of its proteins, with preferential replacements similar to those observed between thermophilic and mesophilic bacteria [6,7,14,15]. We compared the amino acid composition of the predicted proteins of orthologous genes from a variety of Metazoa, including ten annelids of which four were vent species (see methods). Principal component analysis of these compositions showed a dominant axis correlating with GC content of the codons encoding amino acids, and roughly following the trend of genomic GC content (**Figure 5**). Far from being an outlier though, *Alvinella pompejana* has the closest Euclidean distance to the intersection of the two axes. Despite the fact that it has the highest GC content (43%) of the sequenced annelids, its position contrasts strongly with the other vent annelids of lower GC content (∼40%). That amino acid usage tracks GC content has long been noted [48]. In an analysis of bacterial genomes, Kreil & Ouzounis observed a similar trend, but their second principal component neatly distinguished thermophilic from non-thermophilic species [49], whereas we do not see an obvious trend on our 2nd axis.

**Figure 5.**
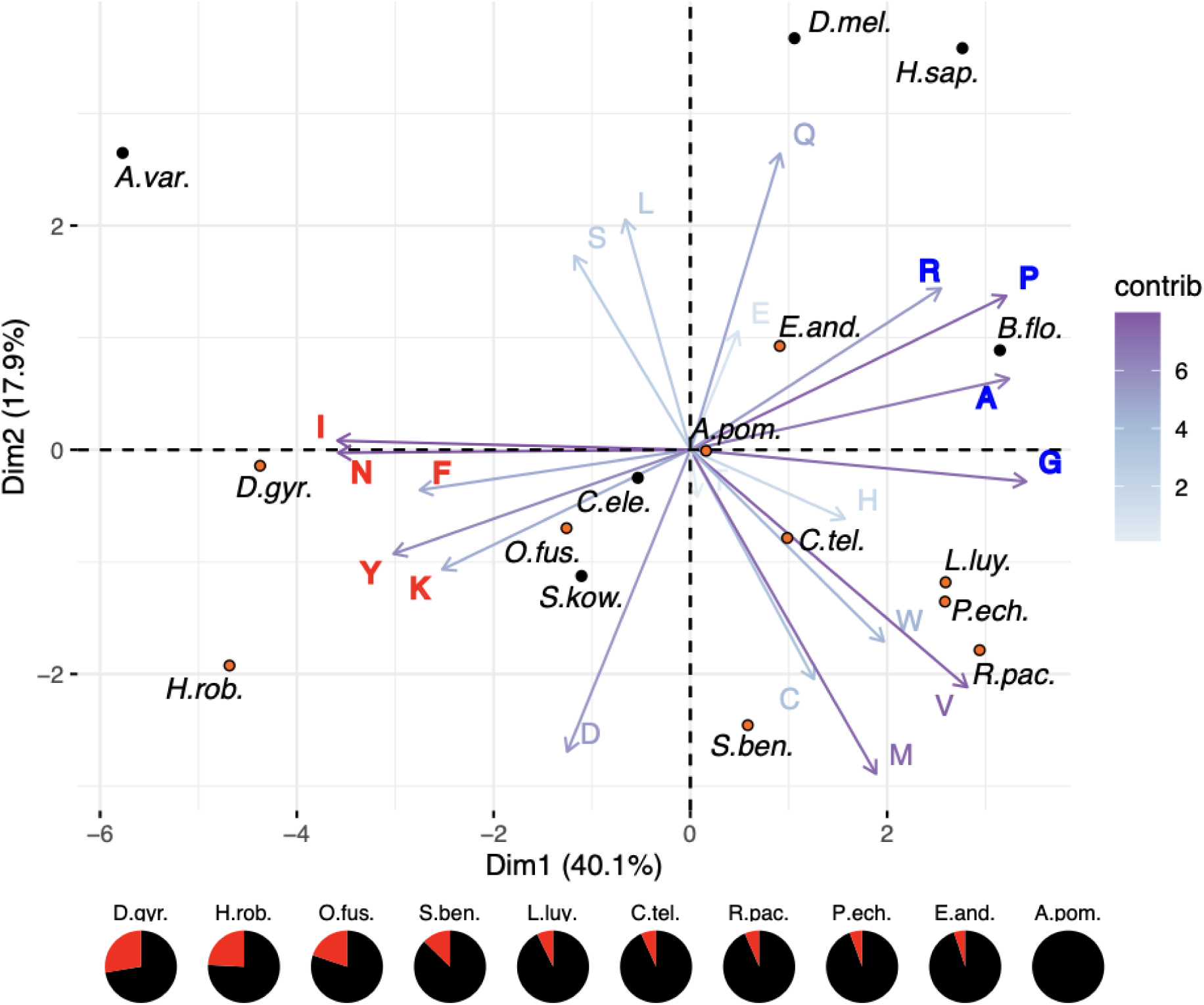
PCA of amino acid composition of various annelids and other Metazoa. Annelid species are indicated by orange dots. Arrows show loading vectors for amino acids (i.e. the extent to which they contribute to the illustrated dimensions) - the darkness (closer to purple) indicates the extent of their contribution. Dimension 1 is dominated by the GC content of codons encoding the amino acids, with low GC (YFINK - red) and high GC (GARP - blue) amino acids at opposite extremes. Genomic GC content as a percentage of the GC content of *Alvinella* (black) for each annelid is shown below the plot, with extra AT content (red) making up the remainder (cf. table S1). Non annelid taxa: A.var *Adineta vag*a; B.flo *Branchiostoma floridae*; C.ele *Caenorhabditis elegans*; D.mel *Drosophila melanogaster*; H.sap Human; S.kow *Saccoglossus kowalevskii*

### Synteny

We used our chromosome scale assembly to examine conserved synteny between *Alvinella* and other Bilateria by equivalencing orthologous genes [50]. Our analysis showed well conserved synteny between *Alvinella* and *Branchiostoma* (**Figure S2**), indicative of good conservation of chromosomes between *Alvinella* and the protostome / deuterostome ancestor, which is not universal in annelids [30].

Our assembly contained 17 segments longer than 1 Mb. An earlier determination of karyotype reported 2n = 32 chromosomes, but noted experimental difficulties [29]. The analysis of the synteny between *Alvinella* and the karyotype of the scallop *Patinopecten yessoensis,* which shows high conservation with inferred ancestral bilaterian chromosomes [51], showed that all of the assembled chromosomes of these species have either a 1:1 equivalence or result from fusions with mixing of the ancestral chromosomes (**Figure 6**).

**Figure 6.**
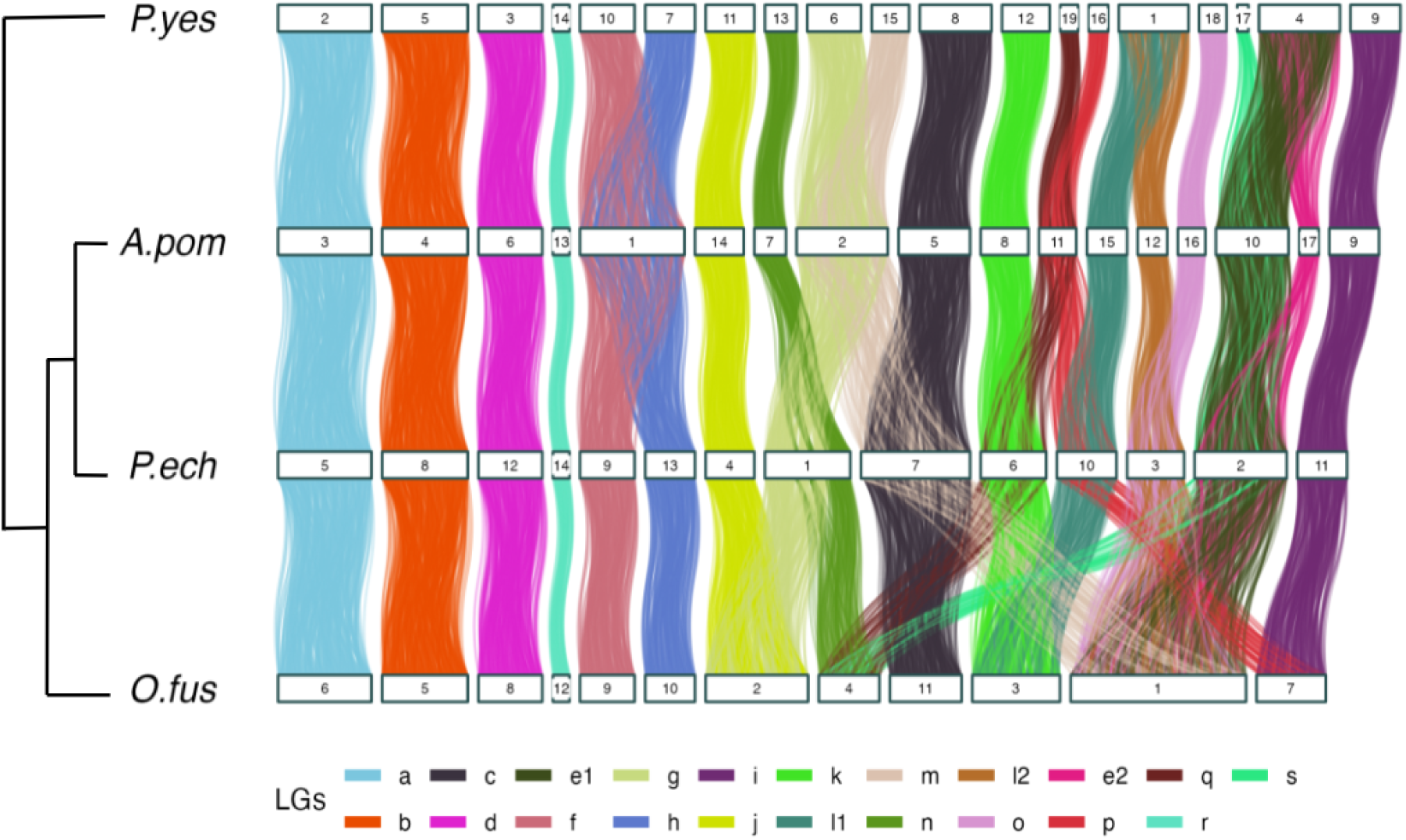
Chord diagram of the inherited karyotypes in annelids. The thermal vent worms *Alvinella* and *Paraescarpia* are compared with the scallop *Patinopecten yessoensis* and the deep branching annelid *Owenia fusiformis*.

### Gene content

#### ANTP-like homeoboxes and other transcription factors

*Alvinella* contains an annelid-like homeobox cluster of around 1Mb in length. In contrast to siboglinids [20], an ANTP gene is present (**Figure 7**). These genes are on the same scaffold as DLX as well as MNX, GBX and EN (the components of the so-called EHGbox), as in *Platynereis dumerilii* [52]. Unlike siboglinid annelids [19], only one copy of Engrailed is present. The parahox genes CDX and GSX are within 100 kb of each other, with PDX more distant, showing some disruption of the likely plesiomorphic state. Interestingly, the transcription factors of the so-called ‘pharyngeal cluster’, a proposed deuterostome synapomorphy [53], including PAX1/9, FOXA, MSXLX and NKX2.1/2.2 are all present within around 250 genes of each other on the same scaffold, adding support to the possibility that this organisation is in fact plesiomorphic for the Bilateria [54]. A variety of NK-like homeodomain proteins are encoded on a further scaffold (EMX, NK2-3/5/6, NK3, MSX, VAX, LBX, TLX, NK1, HMX, HHEX, NOTO, NK6 and NK7) with varying degrees of proximity, suggestive of breakup of a tighter previous degree of clustering, as proposed for the NK-cluster of the ancestral bilaterian [55].

**Figure 7:**
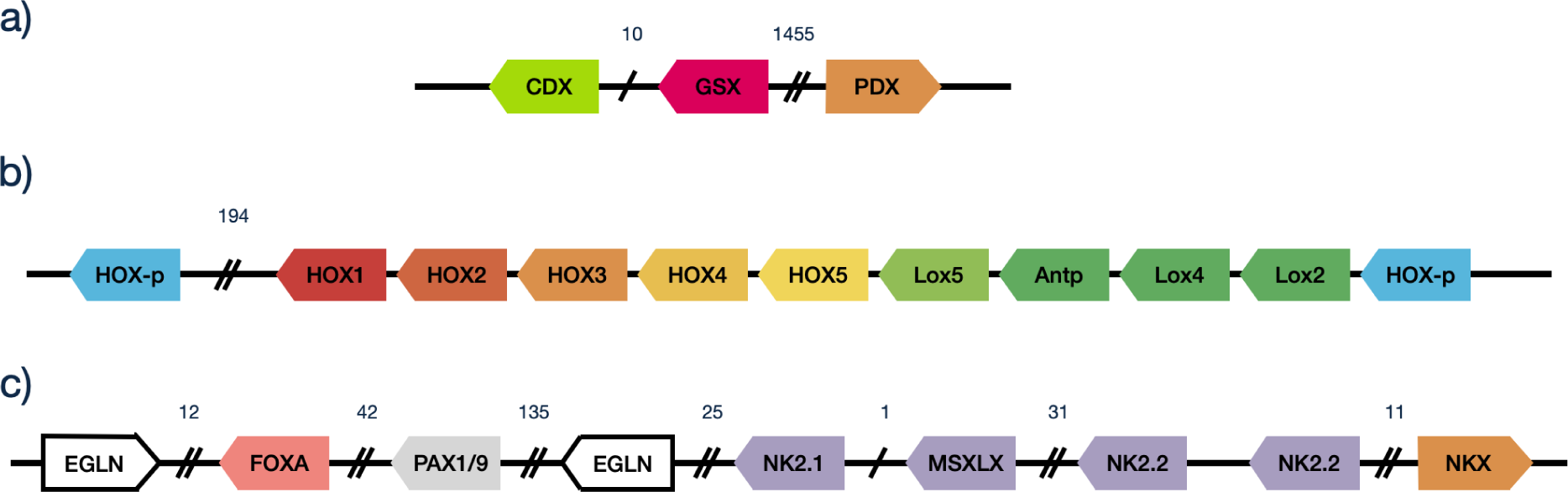
*Alvinella* Parahox, HOX and pharyngeal gene clusters. **a)** parahox cluster **b)** HOX cluster; several likely spurious transcripts are not shown in the posterior part of the main cluster **c)** genes of the deuterostome pharyngeal cluster. MSXLX was not identified in the gene build, but is present in the genome sequence. Numbers above intergenic regions are approximate numbers of intercalated genes.

Among Forkhead transcription factors, we noted the presence of FOXI. Although found in *Capitella* and some other annelids, it is not present in siboglinid genomes [56] and typically absent in protostomes. FOXI is associated with gill structures in deuterostomes, and is a marker for ionocytes [57–59]. Both siboglinids and Alvinellids have extensive gill structures involved in gas transfer, but the shared presence of FOXI in *Alvinella* and deuterostomes hints at the presence of shared ionocyte cell-types lost in other taxa.

#### Collagen

The cuticle of *Alvinella* has been noted to have unusual supercoiled collagen fibrils [60]. Animal genomes encode many collagens, structural proteins composed of multiple repeating triplets of Gly-Xaa-Yaa amino acids. Three collagen chains combine to form a triple helical structure. Thermal stability has been related to the presence of 4-hydroxyproline at the Y position of the repeat [61]. An *Alvinella* interstitial collagen has been highlighted as a protein of unusual thermal stability [62]. We identified this protein (AAC35289.2; 890 residues) as a truncated version of one of our predictions (APscaff0012.263.p1; 1483 residues). We identified 42 predicted *Alvinella* proteins containing hits to the Pfam Collagen HMM (see methods) several of which are extremely long, the longest of which having 8859 amino acids. Although it is difficult to rule out that the repetitive nature of collagen molecules might have led to genomic misassembly artefacts, we detected similarly long proteins in assembled transcriptomes of *P. pandorae* (8689 aa), *P. grasslei* (8413 aa) *P. fijiensis* (7719 aa) (**Table 1**). Notably, no collagens from the recently sequenced siboglinid gene predictions reach a similar length. The earthworm *Eisenia andrei* does include such a collagen, but, as has been noted before in other earthworms, with a much higher proportion of proline at the Xaa position, which is not associated with thermal stability [61,63]. Although we cannot quantify 4-hydroxyproline content from protein sequence alone, the high proline counts at the Yaa position for *Alvinella* and its overall high proline content are consistent with increased thermostability. These trends in collagen proline content run counter to the global amino acid usage that might be expected from the PCA plot presented earlier.

**Table 1:** Annelid collagen-like proteins with the most GXY repeats. Repeats were counted for each match to the pattern shown at the head of the second column. The proportion of repeats with a proline at positions X and Y are shown. P_gra, A_pom, P_pan and P_fij are alvinellids; R_pac, L_luy and P_ech are siboglinids; E_and is an earthworm. P_pan and P_gra are cold adapted annelids, whereas P_fij and A_pom live on vent chimney walls [15].

|  | Gxx**GXY**GxxG | **G-P-Y** % | **G-X-P** % |
| --- | --- | --- | --- |
| P_gra\|DN108_c0_g3_i3-orf | 5722 | 0.386 | 0.264 |
| **A_pom\|APscaff0012.353.p1** | **5698** | **0.372** | **0.333** |
| E_and\|GWHPACBE014886 | 5660 | 0.403 | 0.215 |
| P_pan\|DN152_c0_g2_i2-orf | 5628 | 0.331 | 0.217 |
| P_fij\|DN15_c0_g4_i1-orf | 5012 | 0.332 | 0.326 |
| R_pac\|jg13387.t2 | 840 | 0.133 | 0.362 |
| L_luy\|FUN_004270-T1 | 800 | 0.09 | 0.34 |
| P_ech\|Scaf3834_5.3 | 684 | 0.211 | 0.231 |

#### Tube composition

*Alvinella* builds a stable, watertight, parchment-like tube, enabling it to regulate its thermal microenvironment by drawing in cooler water with up and down body movements. Gaill and Hunt investigated the amino acid composition of these tubes, finding them made of a glycoprotein matrix, in contrast to the chitin/proteoglycan mix of the siboglinid *Riftia* [64], and Vovelle and Gaille reported secretions rich in sulphated glycosaminoglycans [65]. Amino acid composition revealed an abundance of the small amino acids serine, glycine and alanine, in a manner reminiscent of silk fibroins [64,66] and barnacle shell proteins [67]. For all *Alvinella* proteins, we calculated the Euclidean distance to the tube percentages of serine (25%), glycine (20.6%) and alanine (11.8%). The best scoring match was confidently predicted to form an unusually large extended β-sheet structure (**Figure 8**), and although speculative, it is possible that this and similar proteins oligomerize as a fibrillar structural component of the tubes.

**Figure 8:**
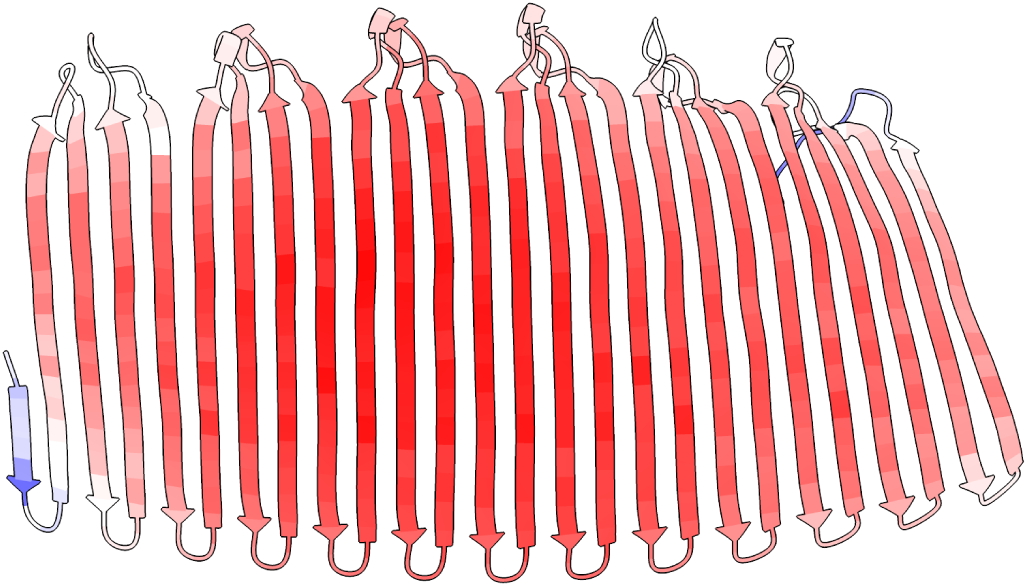
Extended β-sheet structure of the *Alvinella* protein with S,G,A amino acid composition most similar to the reported values for tube composition. The strands are 17 amino acids long. Predicted using Alphafold2, coloured by pLDDT values, red = high, i.e. more confident.

In other annelids, the Tyrosinase gene family has been implicated in modification of tyrosine rich protein families to enable DOPA mediated cross linking of ‘glue’ anchor proteins, and Tyrosinase gene family expansions have been observed [68]. *Alvinella*, however, encodes no matches to the Pfam Tyrosinase domain (see methods).

#### Globins

In common with other annelids [69,70], the *Alvinella* genome encodes a high number of globin genes, predicted to include both intra- and extra-cellular proteins. We detected 2 genomic clusters of globins corresponding to *Alvinella pompejana* specific gene duplications. The first, of 3 members (APscaff0001.1370.p1, APscaff0001.1371.p1, APscaff0001.1372.p1) is predicted to encode extracellular globins, each with the key histidine residues that coordinate the Zn^2+^ ions proposed to bind H_2_S [71]. The second genomic cluster, of 5 adjacent globins, encoded proteins lacking signal peptides which, accordingly, are unlikely to be extracellular. Among our annelid data, they were most closely related to *Eisenia* proteins, with the tardigrade kumaglobin being the closest known structure [72].

Within the large clade of annelid extracellular globin genes we identified further *Alvinella* specific gene duplications, for a total of 11 genes. These *Alvinella* specific duplications are independent of the main expansion of ‘B1’ type globins found in siboglinids [17–19]. A single *Alvinella* gene had a free cysteine similar to ‘B2’ type annelid globins (C+1), while the genomic cluster of 3 members all had an ‘A2’-like free cysteine (C+11), compatible with proposed H_2_S transport (**Figure S3**) [73]. *Alvinella* further encodes 4 linker genes involved in scaffolding the giant extracellular globin complex [74], a number comparable to other annelids, although these again appear to be related by independent duplication and loss events rather than straightforward orthology.

Another unusual globin with dehaloperoxidase activity has been characterised from the terebellid annelid *Amphitrite ornata* [75] and the similarity of an *Alvinella* globin noted [76]. We identified 4 unclustered but paralogous *A. pompejana* genes with a likely orthologous relationship to this gene, but no equivalent siboglinid sequences. In *A. ornata*, it has been speculated that these enzymes detoxify bromoaromatic compounds released by species living in close proximity (*Notomastus lobatus,* a polychaete and *Saccoglossus kowalevskii*, a hemichordate) [77]. In alvinellids, they may have a similar role in detoxifying vent chemicals or bacterial metabolites.

The unrelated oxygen transport protein, hemerythrin, was not detected in the *A. pompejana* proteome, or alvinellid proteins, despite its frequent occurrence in annelids, including siboglinids, and a previous report in *Paralvinella palmiformis* (KY007423.1); [78]. Further analysis of the *P. palmiformis* sequence revealed it to be identical at the protein level to a sequence from the unrelated annelid *Oenone fulgida*, suggesting the possibility of sample contamination and the genuine absence of this family from alvinellids.

#### Hypoxia

The increased numbers of globins observed in annelids, relative to other Metazoa are suggestive of demanding oxygen availability. Some taxa undergoing periodic hypoxia (tardigrades, barnacles and copepods) have lost components of the HIF hypoxic response pathway [79,80]. In *Alvinella* and siboglinids, we detected a full complement of HIF pathway genes (HIF1, HIF1AN, VHL and EGLN). HIF orthologs included the conserved motifs embedding the key proline that is hydroxylated and then bound by VHL prior to destruction [81] (**Figure S4**).

We did, however, uncover other genomic evidence for tolerance of limited oxygen environments, finding that the *Alvinella* genome encodes a pyruvate:NADP+ oxidoreductase (PNO), a marker of eukaryotic anaerobic pyruvate metabolism [82] and a soluble fumarate reductase (similar to OSM1 and FRD1 in *S. cerevisiae*) (supplementary information). These genes were also present in siboglinids and other annelids, as well as some non-chordate bilaterian taxa. In yeast, OSM1 and FRD1 have been implicated in maintaining an oxidising environment under anaerobic conditions [83]. In *Alvinella*, we also detected genes for two octopine dehydrogenase-like proteins (supplementary information). These catalyse opine synthesis, the reductive condensation of pyruvate with arginine (yielding octopine) alanine (alanopine) or glycine (strombine), allowing the anaerobic generation of ATP. This pathway may be advantageous relative to lactate formation as it is osmotically neutral [84]. Octopine dehydrogenases have a sparse phylogenetic distribution, found in cnidarians, molluscs and some annelids. Notably, we could not detect them in siboglinids. Use of rhodoquinone as an electron donor for fumarate is another diagnostic of anaerobic metabolism [84,85]. In animals, its synthesis requires a Co-enzyme Q2 (COQ2) isoform including the ‘e-form’ exon, which is spliced in a mutually exclusive manner with an ‘a-form’ exon [86]. We found both the ‘a-form’ and ‘e-form’ exons of COQ2 adjacent within the *Alvinella* genome (**Figure 9**), and transcript evidence from other alvinellids that mutually exclusive splicing occurs. Finally, we detected a pair of tandemly duplicated alternative oxidase (AOX) genes, which can serve to oxidise ubiquinol in parallel to complex III. In combination, and taken together with the presence of a standard OXPHOS system, these genes are indicators of flexible aerobic mitochondria, using AOX or complex III to oxidize quinones, or even facultatively anaerobic mitochondria that use fumarate as an electron acceptor [85].

**Figure 9:**
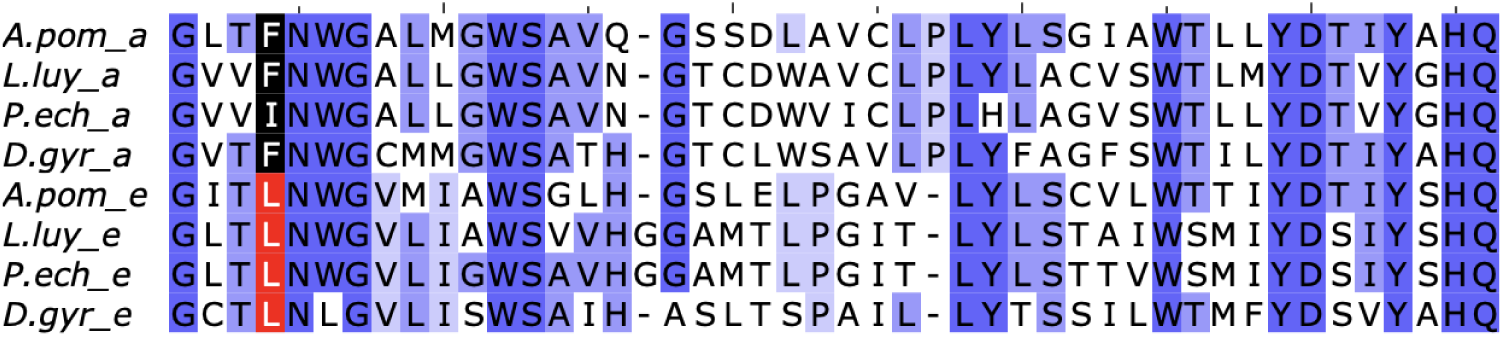
Tandemly duplicated exons in a selection of annelid COQ2 genes. ‘e’-type exons are implicated in rhodoquinone synthesis, and are present in *Alvinella* and siboglinids. Tan and co-workers [86] highlight two residues differentiating these exons - our data suggest that the first of these is most diagnostic (red/black colouring) - the second (third residue from C-terminus, A/S in Tan et al.) does not show a clear pattern with our species sampling. Purple shading reflects the degree of sequence identity for the column. A.pom = *Alvinella pompejana*; L.luy = *Lamellibrachia luymesi*; P.ech = *Paraescarpia echinospica*; D.gyr *= Dimorphilus gyrociliatus*.

#### Kynurenine pathway: present in *Alvinella*, lost in siboglinids

Animal synthesis of the rhodoquinone mentioned above has been shown to be dependent on the kynurenine pathway, via the 3-hydroxyanthranilate (3-HAA) substrate choice by COQ-2 [86–88]. While comparing gene content in *Alvinella* to that of siboglinids, we detected a striking absence in siboglinids (*Riftia, Lamellibrachia* and *Paraescarpia*) of key kynurenine pathway enzymes. We were unable to detect siboglinid orthologs of TDO, KMO, HAAO and QPRT and the associated genes ACMSD, ALDH8A1 and CAT (**Figure 10**). All these missing genes are present in *Alvinella pompejana*, although their distribution in Metazoa as a whole is somewhat patchy. Remaining pathway genes, KYNU and IDO were present in siboglinids, but the overall extent of losses suggests that the pathway is unlikely to function canonically, and its ability, as described, to synthesise 3-HAA questionable. Siboglinids do encode the ‘e’ exon form of COQ2 (**Figure 9**), suggesting that they do use 3-HAA but obtain it *via* another uncharacterized biosynthetic route, perhaps involving a promiscuous enzyme, or from their symbionts. This may limit their capacity to cope with anaerobic conditions, relative to *Alvinella*.

**Figure 10:**
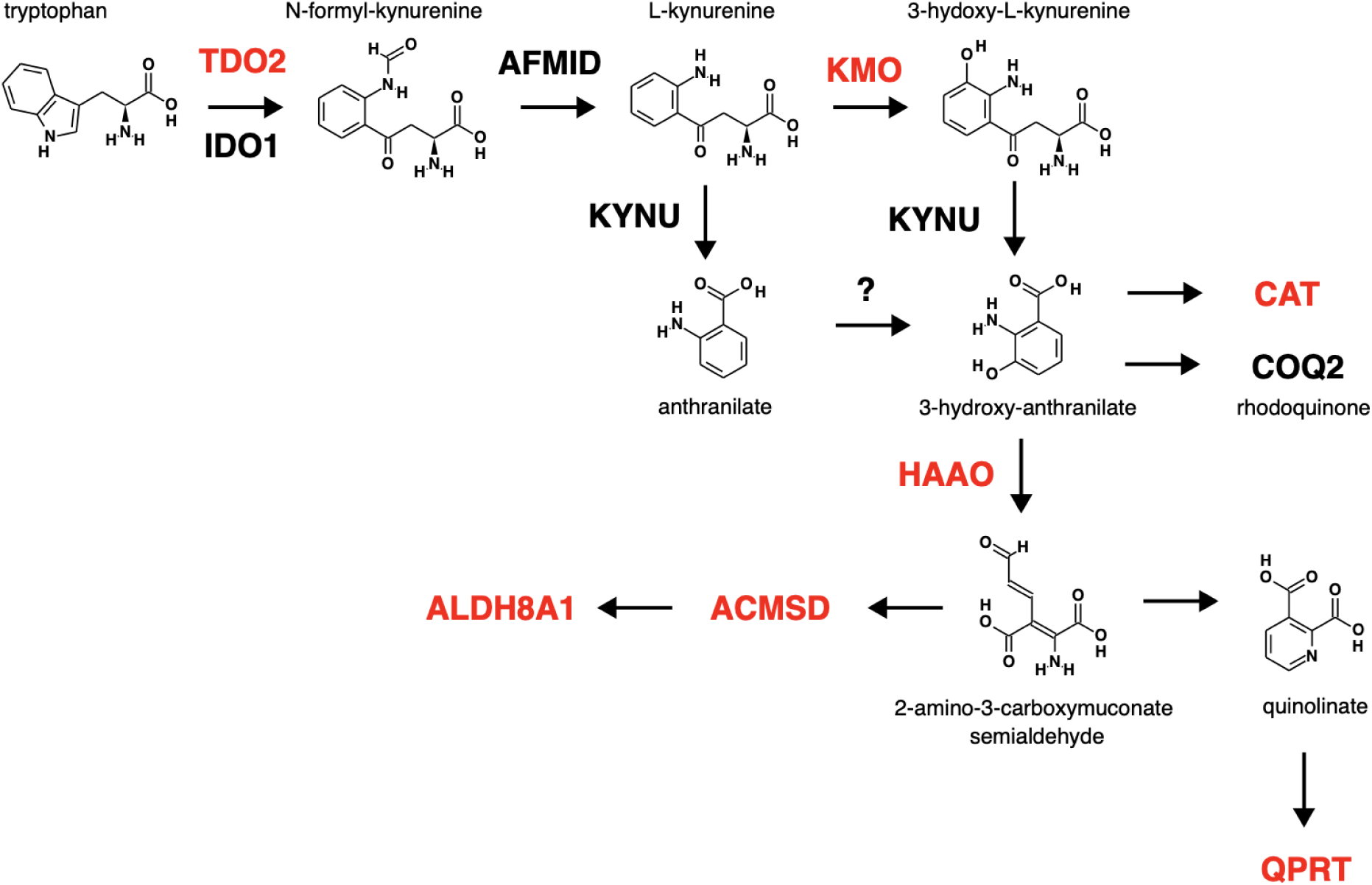
Kynurenine pathway genes. Human gene names are shown in capital letters and those coloured red are present in *Alvinella* & human, but absent from siboglinids. Pathway adapted from Kegg database pathway for tryptophan metabolism (https://www.genome.jp/pathway/map00380).

#### Urea cycle

Enzymes involved in urea excretion were reported absent from an early *Alvinella* transcriptome database [6]. In our genome sequence, we found genes for three out of four urea cycle enzymes: argininosuccinate lyase (ASL), Argininosuccinate synthetase 1 (ASS1) and Arginase (ARG1/2), with the missing exception being Ornithine transcarbamylase (OTC). Vestimentiferan siboglinids, but not the bone-eating *Osedax*, encode all four of these urea cycle enzymes [19,20]. It is notable that *Alvinella* encodes Nitric Oxide synthase genes, enabling a direct arginine-citrulline connection. Although present in other alvinellids and annelids, we found no NO synthase genes in siboglinids. The significance of this is not readily apparent, but conceivably related to use of arginine in octopine synthesis (see above).

#### Absence of light sensitive proteins

The vents that *Alvinella* inhabits are typically several kilometres below the surface, a depth where there is no ambient light source. We did not detect visual opsins in our predicted gene set, consistent with other deep sea annelids [89]. In contrast, the deep sea mussel *Bathymodiolus platifrons* does encode opsin-like genes [90].

We were also unable to detect cryptochrome homologs, light sensitive proteins that regulate the circadian clock. Cryptochromes appear to be absent from many annelid genomes (including the leech *Helobdella* and siboglinids), although they are present in *Capitella* and *Owenia*. Several other circadian clock components were also undetectable in *Alvinella*, including PER, TIMELESS *(sensu tim* in *Drosophila),* CLOCK, and BMAL1 (ARNTL). Although *Alvinella* does not appear to be remarkable in this regard as the distribution of these genes is patchy within Metazoa as a whole, it is notable that they are present in the deep sea mussel *Bathymodiolus azoricus [91]*. Against these absences, it should be noted that tidal rhythms are likely to be relevant to deep sea vent communities e.g. [91,92].

#### An unusual number of carbohydrate sulfo-transferases and esterases

We analysed protein family content using the PANTHER protein database [93], assessing statistically significant enrichment of families in *Alvinella* relative to siboglinids (see methods). Many PANTHER families have related or identical functions, so we then used the family classifications to label genes with Gene Ontology (GO) terms from the ‘molecular function’ namespace and calculated the enrichment of GO terms from the genes encoding the significant set of PANTHER families, relative to the complete *Alvinella* protein coding gene set [94]. (**Table 2**).

**Table 2.** Overrepresented PANTHER families in *Alvinella pompejana. Alvinella* annotations were compared to a single combined proteome of the siboglinids *Riftia*, *Paraescarpia* and *Lamellibrachia*. Significant results are shown (α < 0.01, see methods). Hits to likely reverse transcriptases, and those where the *Alvinella* sequences were predominantly low complexity have been removed (Compare with **Table S8** for raw output)

|  | n | p-val | p-adj | description |
| --- | --- | --- | --- | --- |
| PTHR45953 | 35 | 7E-23 | 3.6E-19 | IDURONATE 2-SULFATASE |
| PTHR15723 | 34 | 2.6E-20 | 8.8E-17 | CARBOHYDRATE SULFOTRANSFERASE 15 |
| PTHR43806:SF11 | 25 | 1.9E-18 | 4.1E-15 | CEREVISIN-RELATED |
| PTHR10704 | 45 | 4.2E-15 | 7.1E-12 | CARBOHYDRATE SULFOTRANSFERASE |
| PTHR47032:SF1 | 31 | 9.7E-13 | 1.2E-09 | UDP-D-XYLOSE:L-FUCOSE<br>ALPHA-1,3-D-XYLOSYLTRANSFERASE-RELATED |
| PTHR34009 | 16 | 4.6E-12 | 5.2E-09 | PROTEIN STAR (Pfam:Methyltransf_21) |
| PTHR12137 | 39 | 1.3E-11 | 1.3E-08 | CARBOHYDRATE SULFOTRANSFERASE |
| PTHR33604:SF3 | 24 | 4.2E-10 | 3.9E-07 | OSJNBA0004B13.7 PROTEIN |
| PTHR10704:SF44 | 12 | 3.2E-09 | 2.7E-06 | PROTEIN, PUTATIVE-RELATED (Pfam:Sulfotransfer_3) |
| PTHR46780:SF9 | 42 | 8.6E-08 | 6.7E-05 | PROTEIN EVA-1 (Pfam:Gal_Lectin) |
| PTHR31511 | 12 | 1.9E-07 | 0.00013 | PROTEIN CBG23764 |
| PTHR11022:SF41 | 9 | 4.2E-07 | 0.00025 | PEPTIDOGLYCAN RECOGNITION PROTEIN 5 |
| PTHR11214:SF314 | 9 | 4.2E-07 | 0.00025 | BETA-1,3-GALACTOSYLTRANSFERASE 4 |
| PTHR32385:SF20 | 12 | 7.9E-07 | 0.00045 | MANNOSYL PHOSPHORYLINOSITOL CERAMIDE SYNTHASE<br>CSH1-RELATED |
| PTHR25462 | 25 | 8.5E-07 | 0.00046 | BONUS, ISOFORM C-RELATED |
| PTHR24028 | 35 | 7.5E-06 | 0.0033 | CADHERIN-87A |
| PTHR47327:SF2 | 7 | 1.1E-05 | 0.0047 | FI18240P1-RELATED (Pfam:PAN) |
| PTHR21461 | 10 | 1.3E-05 | 0.0053 | UNCHARACTERIZED (Pfam:Glyco_transf_92) |

This PANTHER family analysis showed that both carbohydrate sulfotransferases and esterases (Iduronate 2-sulfatase) were over-represented, along with a β-1,3-galactosyltransferase family. This suggests a complex but unexplored sulfate modified carbohydrate chemistry, and it is of note that sulfated glycosaminoglycans are important in tube formation [65]. Sulfotransferases also represent an expanded gene family (27 genes) in vent shrimps together with sulfatases, some of which are positively-selected and overexpressed [95]. It seems likely, therefore, that these molecules aid *Alvinella* in its thermal environment or in bacterial symbioses.

Given these hits, enriched GO terms relative to the *Alvinella* protein content as a whole included ‘sulfotransferase activity’, ‘serine-type peptidase activity’ as well as ‘mannosyltransferase activity’ and ‘glycosyltransferase activity’. Notably absent from the results of this analysis were heat shock proteins, for which gene family expansions have been reported in the deep sea vent mussel *Bathymodiolus platifrons* (HSP70) [90], compared to constitutive high expression in *Paralvinella sulfincola* [96].

## Discussion

A variety of deep-sea vent invertebrates have recently had their genomes sequenced, including the mussel *Bathymodiolus platifrons* [90], the clam *Archivesica marissinica* [97], the scaly-footed snail *Chrysomallon squamiferum* [98], the snail *Gigantopelta aegis* [99]; among annelids, the scale worm *Branchipolynoe longqiensis,* several siboglinids and the alvinellid *Paralvinella palmiformis* [25]; among cnidarians, the deep sea anemones *Alvinactis idsseensis* sp. Nov. [100] and *Actinernus* sp. [101], but their relative inaccessibility means that they are sparsely represented in the literature, and global studies on adaptations to their unique environmental challenges not common.

The challenge of living on deep sea vents begins with colonization, and genome sequences will facilitate future work using population genomics to study community genetic connectivity [102]. Incipient species are common in the deep sea [103–105], and indeed, we find that *A. pompejana* shows distinct populations on either side of the equator. These two lineages showed classic speciation associated features, with heterogeneous divergence across the genome [106], and asymmetric introgression (e.g. [107,108] from the North EPR into the South EPR. This introgression was detected in two samples located thousands of kilometres away from the contact zone [27]. Such long distance introgression may be more frequent along mid-oceanic ridges due to the emergence of physical dispersal barriers following tectonic rearrangements [109].

The best-studied metazoan species in terms of adaptation to high temperatures associated with some vent types is the Pompeii worm *Alvinella pompejana*. Although *Alvinella’s* protein amino acid composition can be distinguished from other species [7,14,15], in a manner that might correlate with ambient temperature, given the paucity of such studies across the full diversity of animals, and the corresponding dearth of comparators, it is not clear that definitive evidence has been produced. The analysis presented here instead suggests that *Alvinella* bulk protein amino acid composition is not unusual (**Figure 5**), but this does not preclude the possibility that some proteins, or groups of proteins sharing particular expression patterns, may be particularly thermotolerant (as suggested by composition of collagen triplets discussed above for instance). Large-scale protein thermal stability assays [110] coupled with more stratified protein groupings may help clarify these observations.

To adapt to temporally high temperatures, *Alvinella* has developed a thick, collagen-enriched integument with particularly high thermal stability [62] and builds a stable tube that may limit exposure to scalding water [111]. We were able to identify unusually long collagens, showing sequence features indicative of thermal stability. We also propose potential tube component proteins, and suggest that extensive duplication of genes involved in sulfate modified carbohydrate chemistry may be related to tube construction.

By comparing *Alvinella* to other deep sea vent worms, the siboglinids, we have highlighted similarities and differences in deep sea survival strategies. We find that both siboglinids and *Alvinella pompejana* encode a rich variety of globins and secreted globins, and that, to an extent, these reflect independent diversifications. We find that *Alvinella*, like the siboglinids and some other but not all deep sea species, is characterized by the secondary absence of light sensitive opsins and cryptochromes. Both *Alvinella* and the siboglinids contain enzymes that are characteristic of facultatively anaerobic mitochondria (PNOR and the exon required for COQ2 rhodoquinone synthesis), but that in addition to this, *Alvinella* encodes proteins for octopine synthesis, NO synthesis and a complete kynurenine pathway. These adaptations to anaerobic environments are not unique to deep sea annelids, and are likely inherited from a common annelid or even bilaterian ancestor [84,112]. Differential gene expression data for *Alvinella* are not available, but the hot vent annelid *Paralvinella sulfincola* shows a reduction in respiratory complex I protein expression at higher temperatures, consistent with a move away from oxidative phosphorylation and increased dependence on anaerobic metabolism, possibly to reduce reactive oxygen species formation [96].

While our observations have highlighted adaptations to the deep sea, the overall impression is that the *Alvinella pompejana* genome has been evolving in a relatively conservative manner. Representative genomes of closely related taxa from both deep sea and less extreme marine environments will help further pin down its specific adaptations.

## Methods

### Sample collection, DNA extraction, sequencing and assembly

*Alvinella* specimens were collected from the top of a chimney at the vent site and the snowball of the Bio9 site at 9°50N/EPR using the telemanipulated arm of the HOV *Nautile* during the French oceanographic cruise Mescal2012 (chief scientist: N. LeBris). Animals were brought back to the surface in an insulated basket and immediately flash-frozen after dissection in liquid nitrogen. Ultra-high molecular weight genomic DNA (gDNA) was extracted following the Bionano Genomics IrysPrep agar-based, animal tissue protocol (Catalogue # 80002) from *A. pompejana* gill material. Long-read PacBio sequencing and short-read Illumina sequencing were performed at the Genome Centre of the University of California Berkeley using PacBio Sequel II and Illumina Novaseq platforms. An initial assembly was produced using wtdbg2 [113].

### Scaffolding to chromosome level

Additional frozen *A. pompejana* samples corresponding to the anterior part of the worm (head + gills: 10 individuals) were used for further library preparation, sequencing and scaffolding were performed commercially by Dovetail Genomics. Chicago and Dovetail HiC libraries were prepared as described previously [114,115]. The libraries were sequenced on an Illumina HiSeq X to produce 2×150 bp paired end reads. The input *de novo* assembly, Chicago library reads, and Dovetail HiC library reads were used as input data for HiRise, a software pipeline designed specifically for using proximity ligation data to scaffold genome assemblies [114]. An iterative analysis was conducted. First, Chicago library sequences were aligned to the draft input assembly using a modified SNAP read mapper (http://snap.cs.berkeley.edu). The separations of Chicago read pairs mapped within draft scaffolds were analyzed by HiRise to produce a likelihood model for genomic distance between read pairs, and the model was used to identify and break putative misjoins, to score prospective joins, and make joins above a threshold. After aligning and scaffolding Chicago data, Dovetail HiC library sequences were aligned and scaffolded following the same method.

### Gene model predictions and genome annotation

A de-novo repeats library was constructed using RepeatModeler-2.0.1 [116] and the assembly was masked for repeats using RepeatMasker-4.1.0 (http://www.repeatmasker.org/). An intermediate masked genome, with low complexity parts left unmasked, was generated for the purpose of gene predictions. Gene models were generated by using a combination of de-novo, homology-based and transcriptome-based approaches. Three bulk RNA-seq samples were mapped to our reference genome using STAR-2.7.1a [117] and transcriptomes were assembled using StringTie-2.1.0 [118]. A de-novo transcriptome was also constructed using Trinity v2.15.0 [119] and clustered using TGICL [120]. It was used in the PASA pipeline [121] to generate a set of transcripts based gene models. Exonerate-2.4.0 was used to generate a set of protein-based spliced alignments of previously published Alvinellid transcriptomes [15]. These two sets of spliced alignments were used to generate a set of hints to train Augustus-3.2.3 [122] as described in [123]. Augustus generated a set of 17,245 gene models. A final set of 20,098 gene structures was automatically computed as the consensus of all generated evidence using EVidenceModeler-1.1.1 [124]. The PASA pipeline was finally used to generate a set of updated gene models.

### Functional annotation of predicted proteins

The set of predicted protein sequences was annotated against publicly available databases. The hmmsearch function from HMMER-3.3 [125] was used to retrieve the Pfam protein domains [126] and the PANTHER families [93]. The presence of a signal peptide in the protein sequences was predicted by using signalP-5.0b [127]. We used diamond-0.9.22 [128] to find the orthologs of our predicted proteins against an instance of NCBI non redundant protein set downloaded in October 2020 [129]. Finally, the orthologs in the Kyoto Encyclopedia of Genes and Genomes (KEGG) were assigned using the KEGG automatic annotation server blastKOALA [130].

### Mitochondrial genome assembly and annotation

We assembled a complete and circular mitochondrial genome from a set of short Illumina reads using the toolkit GetOrganelle v1.7.7.0 [131]. The toolkit was used as recommended by the authors with background dependencies on Bowtie2 v2.4.1 [132], SPAdes v3.13.1 [133] and Blast+ v2.5.0 [134]. The correctness of GetOrganelle’s output was assessed by visualising the assembly graph using Bandage v0.8.1 [135]. Annotation of protein coding, rRNA and tRNA genes was achieved by submitting the assembled genome to the MITOS web server [136].

### Whole genome resequencing

In addition to the reference genome, we included whole genome resequencing (WGS) data from six samples, four from the EPR North (9°50N) and two from the EPR South (18°50S), to explore the genome-wide diversity in *A. pompejana*. Individual DNAs were extracted from tissue samples using the CTAB protocol [137,138]. WGS libraries were constructed using Illumina Nextera kits. The six libraries were then standardised by DNA shearing to obtain an insert size of 300 to 600 bp using sonication, and pooled in equimolar proportion. The pooled libraries were paired-end sequenced (2 x 125 bp) on MiSeq, targeting a coverage of 15X per sample, at the Genomer platform of Roscoff Biological Station. The data were analysed following the pipeline developed in [139] for a mollusk (*Littorina saxatilis*). In brief, low quality reads (<15 phred score) and adapter sequences were removed from the raw WGS data using ‘fastp’ v0.23.1 (Chen et al., 2018). Cleaned WGS reads were mapped to the *A. pompejana* reference genome produced in this study using ‘bwa-mem’ v0.7.17 [140]. Duplicated reads in the alignment file were marked using ‘samtools’ v1.16 [141]. Mapped reads were processed using ‘mpileup’ variant caller in bcftools v1.16 [141] to extract Single Nucleotide Polymorphism (SNP) data in variant call format (VCF). Insertions and Deletions (Indels), as well as SNPs falling in 5 bp around the indels were filtered out using bcftools v1.16. Finally, only SNPs with phred quality score of 30, a minimum depth of 5, a maximum depth of 50, and no missing data were included for the downstream analyses using the software vcftools v0.1.16 [142]. This dataset was used to explore the variation across the 6 *A. pompejana* samples using PCA analyses with the R package adegenet [143]. Local PCA analyses were then conducted using a sliding windows approach with windows of 100kb to explore the variation of population structure along the genome. Finally, phylogenetic analyses were conducted based on the polymorphism data of genomic regions with contrasted PCA patterns with the R package phangorn [144].

### Assessment of putative bacterial contaminations in annelid genomes

We used diamond-0.9.22 [128] to query the predicted protein sequences of selected annelids against NCBI non redundant protein set downloaded in october 2020 [129]. The predicted proteomes of *Alvinella pompejana* (this study), *Capitella teleta*, *Helobdella robusta* [145], *Lamellibrachia luymesi* [17], *Paraescarpia echinospica [18]* and *Riftia pachyptila* [19] were annotated for putative bacterial contamination. We filtered out every hit to species in the query which were already integrated in the database (here “*Capitella teleta*” and “*Helobdella robusta*”) as these species would hamper the proper identification of contaminating protein sequences. We set the significance threshold to e-value < 1.e-20 and the proteins retained as contaminants are those sequences with no significant hit in “Metazoan” and at least 1 significant hit in “Bacteria”. The sequence identifiers of the putative bacterial contaminations are provided in supplementary material, and the statistics are summarised in supplementary table S7.

### Amino acid composition

For each genome pair, we calculated reciprocal best hits for predicted proteins using ssearch36 from the FASTA package [146]. For each set of orthologous pairs, we calculated an average amino acid use for each species. Then for each species, we averaged amino acid usage over all species pair averages. The resulting matrix of average amino acid usage across species was used as input for the ‘prcomp’ PCA function in R and visualised using the ‘fviz_pca_biplot’ function from the ‘factoextra’ library. Similar results along the first PCA were obtained using non-redundant counting of proteins (i.e. no averaging of averages) or using counts taken from conserved columns of protein sequence alignments (not shown).

### Annelid phylogeny

We used OMA [147] to compute the orthologs of predicted proteins of annelids for which the complete genomes were available (listed supplementary table S1) and the outgroup species *Crassostrea gigas*, *Lingula anatina* and *Phoronis australis*. All included datasets (**Table S5**) were pre-processed to feature only one protein per gene. We selected the 301 single copy orthologs (100278 amino acids) which included all annelids and at least two outgroup species to make multiple alignments using mafft-v7.309 [148]. Gap containing regions were removed using trimAl v1.4.rev15 [149] with the options ‘-gt 1’, and the alignments concatenated. The phylogeny was computed with phylobayes 4.1c using 4 chains with the CAT+GTR model [36,150].

### Synteny analyses

The orthologous protein sequences *Alvinella pompejana* shared with *Paraescarpia*, *Owenia* and *Patinopecten yessoensis,* were inferred from pairwise reciprocal best hits using diamond-0.9.22 [128] through the script rbhXpress [151]. These reciprocal best hits were displayed on Oxford grids using the R package macrosyntR [152]. Single copy orthologs were extracted from a run of Orthofinder [153]. macrosyntR was then used to draw the chord diagram.

### Assembly of DNA likely originating from symbiont

We mapped Illumina paired-end reads from a previous sequencing campaign [24] to our reference assembly using the bwa-mem algorithm of bwa-0.7.17-r1188 [140]. Using samtools-1.15.1, we selected the unmapped but paired reads (-f option requiring samflags 13) [154]. We quality-filter and trim the selected reads using cutadapt [155]. These sequences, not contained in the *Alvinella* assembly, were assembled using metaSPAdes [156] and quality of the produced assembly was evaluated using Quast [157]. Using metabat2 [158], we further binned the contigs into 14 bins of varying quality, assessed using checkM [159].

### Sequence analysis

General sequence database searching was performed with phmmer from the hmmer package [125]. When these produced hits that were clearly distinguishable from other sequence subfamilies, this was taken as evidence of orthology. In other cases, phylogenies were constructed using IQ-TREE [160] following sequence alignment to hidden Markov models (HMM) of the key PFAM representative within the protein of interest [126] and subsequent trimming, as described in [54]*. Globins:* globins were identified by matches to the Pfam Globin HMM. Signal peptides were predicted on these proteins using signalp 5.0 [127]. A phylogenetic tree was inferred, cross-referenced with the signal peptide predictions and the monophyletic clade of annelid secreted globins extracted. *Absence of opsins:* predicted sequences were searched using a hidden Markov model (HMM) constructed from opsin-like protein sequences. To be considered opsins, we required that database hits included the conserved Schiff-base Lysine residue. Searches against the Pfam database used gathering thresholds as cutoffs (‘--cut_ga’ in hmmsearch). *Absence of Octopine dehydrogenases in siboglinids:* no significant hits were found with the Pfam Octopine_DH HMM in the predicted protein sets of *Riftia*, *Lamellibrachia* and *Paraescarpia*. *Absence of NO synthases:* no significant hits were found with the Pfam NO_synthase HMM in the predicted protein sets of *Riftia*, *Lamellibrachia* and *Paraescarpia*. Collagen containing proteins were detected as hits to the Collagen HMM. *Absence of Tyrosinases in Alvinella:* no significant hits to the Pfam Tyrosinase HMM.

### PANTHER family analysis

Proteome sets were searched against the PANTHER 17.0 database, using the supplied pantherScore2.2.pl script with the -B (display best hit), -n (display family and subfamily) and -s (use hmmsearch) options [93]. Enrichment of each PANTHER model was calculated by comparing counts in *Alvinella pompejana* to the summed counts of the siboglinids, using a Χ^2^ test. P-values were corrected for multiple comparisons using the Benjamini/Hochberg FDR with an alpha of 0.01. *Alvinella* genes with matches to significantly scoring PANTHERs were extracted as a studyset and their associated GO terms were tested for enrichment relative to the GO terms for all *Alvinella* proteins (population) using the ‘ontologizer’ tool [94].

## Supporting information

supplementary information including extra figures and some sequences

## Acknowledgements

We gratefully acknowledge support from the Human Frontiers Science Project Organization grant RGP0028/2018. RRC thanks Max Telford for helpful discussions. Thanks to Aldrich Hezekiah for the drawing of *Alvinella* in Figure 1.

## Data availability

*Alvinella* genome, protein set, GFF annotation and metagenome protein predictions are available at: doi:10.5281/zenodo.11241339

## Supplementary Figures

**Figure S1:**
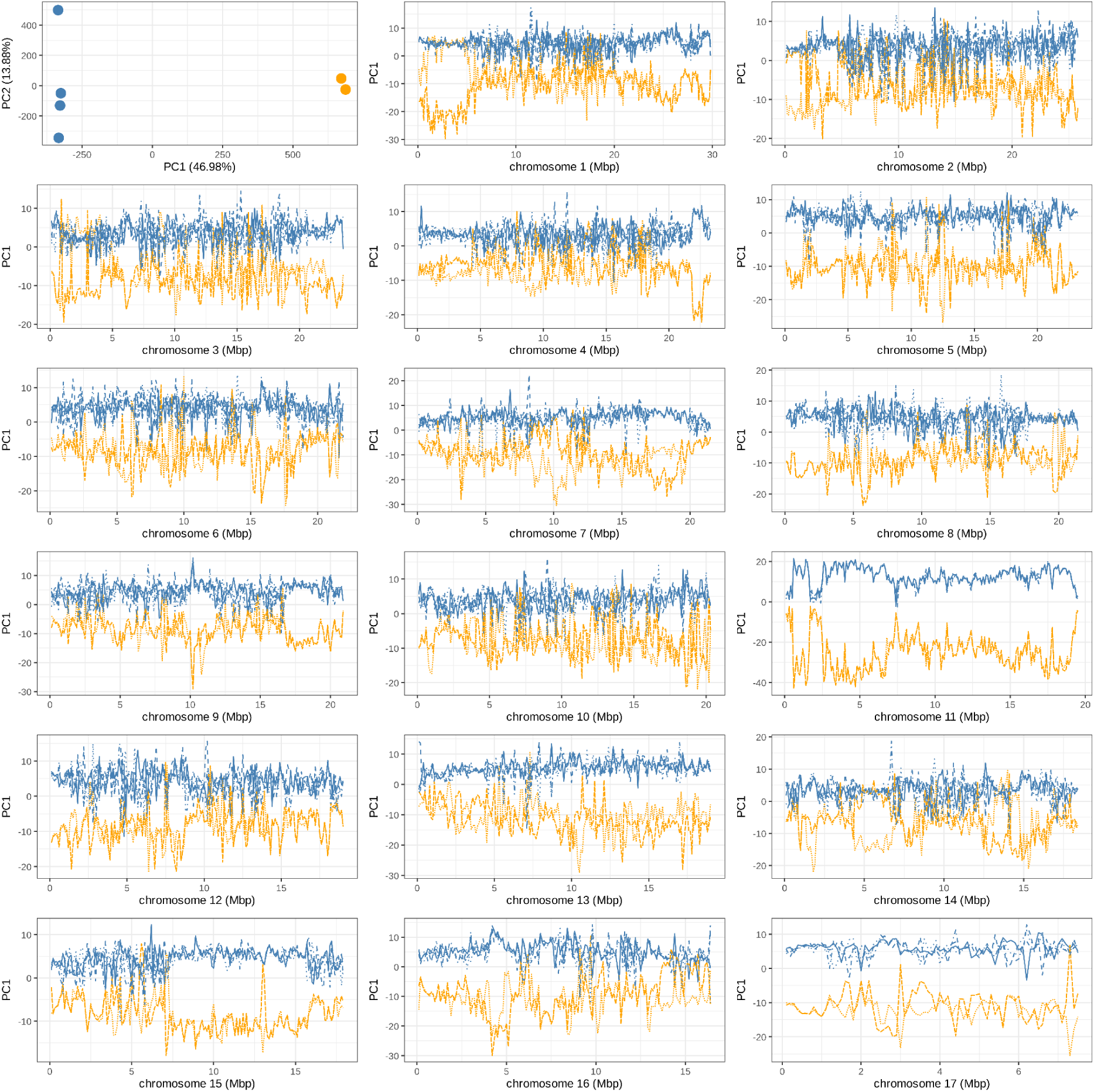
Principal Component Analyses (PCA) performed on the six *A. pompejana* samples genotyped at 3.33M SNPs. The graph on the top left shows the two first axes of PCA computed on the overall dataset. Most of the variation is captured by PC1 (PC1 = 46.98%; PC2 = 13.88%) and distinguishes the two EPR South samples (in orange) from the four EPR north samples (blue). Detailed analyses of genomics regions showing contrasted patterns of divergence between the six samples of *A. pompejana* in a. chromosome 1 and b. chromosome 11. In each subplot, the top graphs show phylogenetic trees computed from the polymorphism data extracted from 1Mbp to 1.5Mbp windows defined from the patterns of divergence observed in the local PCA plot at the bottom. Dotted lines show the limit of the genomics regions that were used to infer the phylogenetic trees. On chromosome 1 (a.), the phylogenies showed, in order of appearance, region of introgression where one EPR south samples is located at mid distance from the EPR north samples and the other EPR south sample, a region of introgression where one EPR south sample clustered perfectly with the EPR north samples, a region of weak divergence where all samples were similarly distant, and a region of high divergence where individuals on either side of the equator were clearly separated. On chromosome 11 (b.), all phylogenetic trees showed clear North – South EPR separation, with the second and last trees also showing very small terminal branches expected from low genetic diversity within each population

**Figure S2:**
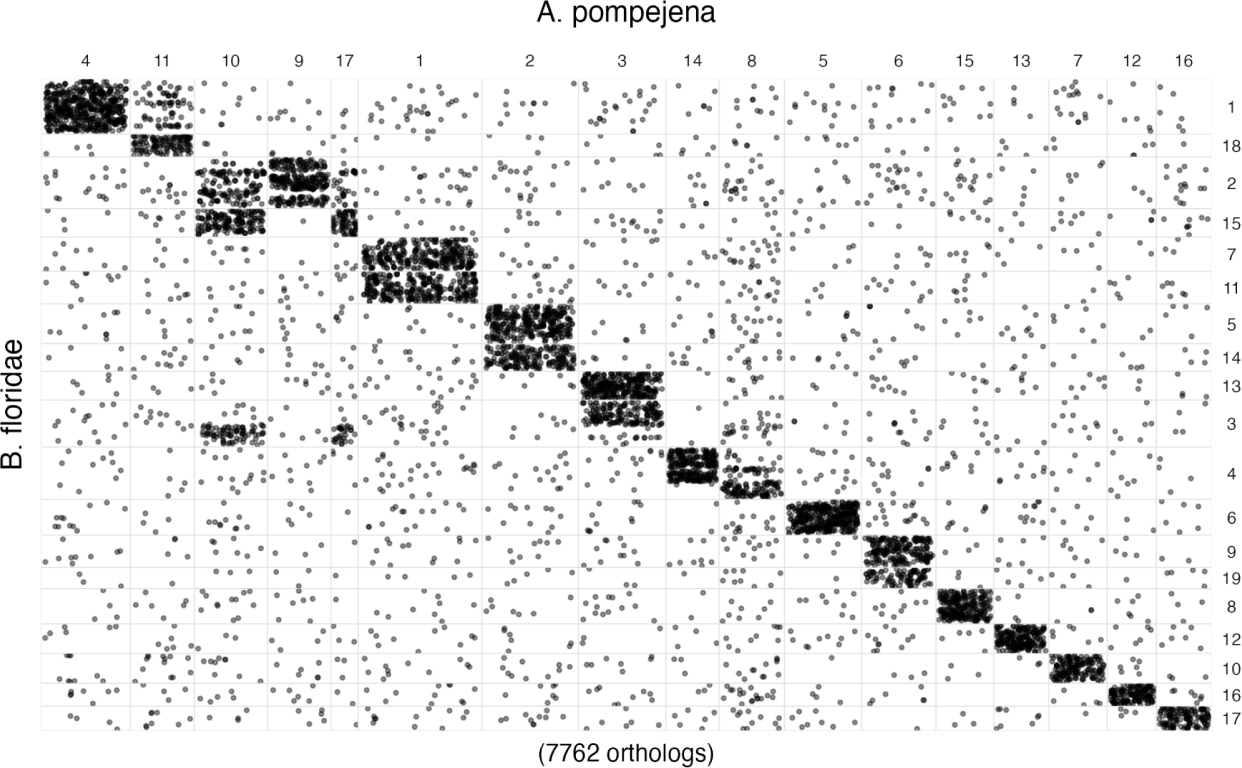
Oxford Grid showing equivalence of *Alvinella* and *Branchiostoma floridae* chromosomal segments.

**Figure S3: Phylogeny of extracellular globins.** *Alvinella* sequence identifiers are in orange.

**Figure S4:**
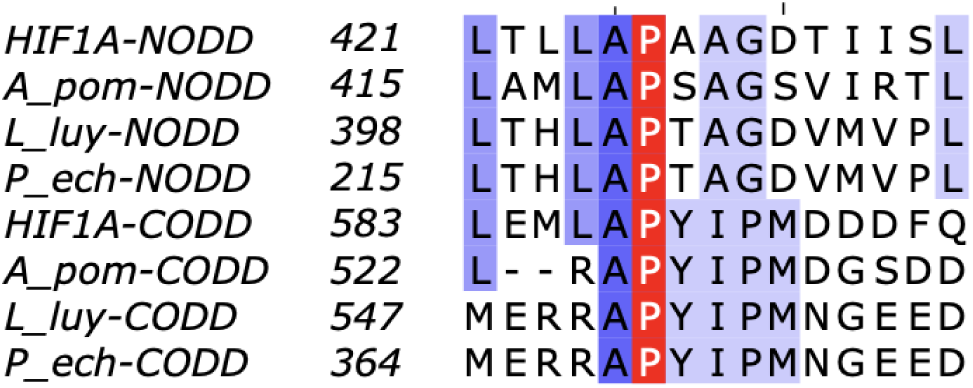
HIF1A N- and C-terminal Oxygen Dependent Degradation motifs are conserved in *Alvinella* and siboglinids. The conserved proline hydroxylation target is shown in red. The human HIF1A sequences are shown with *Alvinella* (A_pom) and the siboglinids (*Lamellibrachia* L_luy, *Paraescarpia* P_ech).

## References

1. Ravaux J, Hamel G, Zbinden M, Tasiemski AA, Boutet I, Léger N, Tanguy A, Jollivet D, Shillito B: Thermal limit for metazoan life in question: in vivo heat tolerance of the Pompeii worm. PLoS One 2013, 8:e64074.

2. Girguis PR, Lee RW: Thermal preference and tolerance of alvinellids. Science 2006, 312:231.

3. Freeman MT, Czenze ZJ, Schoeman K, McKechnie AE: Extreme hyperthermia tolerance in the world’s most abundant wild bird. Sci Rep 2020, 10:13098.

4. Freeman MT, Czenze ZJ, Schoeman K, McKechnie AE: Adaptive variation in the upper limits of avian body temperature. Proc Natl Acad Sci U S A 2022, 119:e2116645119.

5. Bates AE, Lee RW, Tunnicliffe V, Lamare MD: Deep-sea hydrothermal vent animals seek cool fluids in a highly variable thermal environment. Nat Commun 2010, 1:14.

6. Gagnière N, Jollivet D, Boutet I, Brélivet Y, Busso D, Da Silva C, Gaill F, Higuet D, Hourdez S, Knoops B, et al.: Insights into metazoan evolution from Alvinella pompejana cDNAs. BMC Genomics 2010, 11:634.

7. Holder T, Basquin C, Ebert J, Randel N, Jollivet D, Conti E, Jékely G, Bono F: Deep transcriptome-sequencing and proteome analysis of the hydrothermal vent annelid Alvinella pompejana identifies the CvP-bias as a robust measure of eukaryotic thermostability. Biol Direct 2013, 8:2.

8. Shin DS, Didonato M, Barondeau DP, Hura GL, Hitomi C, Berglund JA, Getzoff ED, Cary SC, Tainer JA: Superoxide dismutase from the eukaryotic thermophile Alvinella pompejana: structures, stability, mechanism, and insights into amyotrophic lateral sclerosis. J Mol Biol 2009, 385:1534–1555.

9. Zhang Q, Balourdas D-I, Baron B, Senitzki A, Haran TE, Wiman KG, Soussi T, Joerger AC: Evolutionary history of the p53 family DNA-binding domain: insights from an Alvinella pompejana homolog. Cell Death Dis 2022, 13:214.

10. Wijckmans E, Nys M, Debaveye S, Brams M, Pardon E, Willegems K, Bertrand D, Steyaert J, Efremov R, Ulens C: Functional and Biochemical Characterization of Alvinella pompejana Cys-Loop Receptor Homologues. PLoS One 2016, 11:e0151183.

11. De Gieter S, Gallagher CI, Wijckmans E, Pasini D, Ulens C, Efremov RG: Sterol derivative binding to the orthosteric site causes conformational changes in an invertebrate Cys-loop receptor. Elife 2023, 12.

12. Longo MA, Roy S, Chen Y, Tomaszowski K-H, Arvai AS, Pepper JT, Boisvert RA, Kunnimalaiyaan S, Keshvani C, Schild D, et al.: RAD51C-XRCC3 structure and cancer patient mutations define DNA replication roles. Nat Commun 2023, 14:4445.

13. Chinnam NB, Thapar R, Arvai AS, Sarker AH, Soll JM, Paul T, Syed A, Rosenberg DJ, Hammel M, Bacolla A, et al.: ASCC1 structures and bioinformatics reveal a novel Helix-Clasp-Helix RNA-binding motif linked to a two-histidine phosphodiesterase. J Biol Chem 2024,

14. Jollivet D, Mary J, Gagnière N, Tanguy A, Fontanillas E, Boutet I, Hourdez S, Segurens B, Weissenbach J, Poch O, et al.: Proteome adaptation to high temperatures in the ectothermic hydrothermal vent Pompeii worm. PLoS One 2012, 7:e31150.

15. Fontanillas E, Galzitskaya OV, Lecompte O, Lobanov MY, Tanguy A, Mary J, Girguis PR, Hourdez S, Jollivet D: Proteome Evolution of Deep-Sea Hydrothermal Vent Alvinellid Polychaetes Supports the Ancestry of Thermophily and Subsequent Adaptation to Cold in Some Lineages. Genome Biol Evol 2017, 9:279–296.

16. Stiller J, Tilic E, Rousset V, Pleijel F, Rouse GW: Spaghetti to a Tree: A Robust Phylogeny for Terebelliformia (Annelida) Based on Transcriptomes, Molecular and Morphological Data. Biology 2020, 9.

17. Li Y, Tassia MG, Waits DS, Bogantes VE, David KT, Halanych KM: Genomic adaptations to chemosymbiosis in the deep-sea seep-dwelling tubeworm Lamellibrachia luymesi. BMC Biol 2019, 17:91.

18. Sun Y, Sun J, Yang Y, Lan Y, Ip JC-H, Wong WC, Kwan YH, Zhang Y, Han Z, Qiu J-W, et al.: Genomic Signatures Supporting the Symbiosis and Formation of Chitinous Tube in the Deep-Sea Tubeworm Paraescarpia echinospica. Mol Biol Evol 2021, 38:4116–4134.

19. de Oliveira AL, Mitchell J, Girguis P, Bright M: Novel Insights on Obligate Symbiont Lifestyle and Adaptation to Chemosynthetic Environment as Revealed by the Giant Tubeworm Genome. Mol Biol Evol 2022, 39.

20. Moggioli G, Panossian B, Sun Y, Thiel D, Martín-Zamora FM, Tran M, Clifford AM, Goffredi SK, Rimskaya-Korsakova N, Jékely G, et al.: Distinct genomic routes underlie transitions to specialised symbiotic lifestyles in deep-sea annelid worms. Nat Commun 2023, 14:2814.

21. Saulnier-Michel C, Gaill F, Hily A, Alberic P, Cosson-Mannevy MA: Structure and functions of the digestive tract of Alvinella pompejana, a hydrothermal vent polychaete. Can J Zool 1990, 68:722–732.

22. Dubilier N, Bergin C, Lott C: Symbiotic diversity in marine animals: the art of harnessing chemosynthesis. Nat Rev Microbiol 2008, 6:725–740.

23. Weigert A, Bleidorn C: Current status of annelid phylogeny. Org Divers Evol 2016, 16:345–362.

24. Thomas-Bulle C, Bertrand D, Nagarajan N, Copley RR, Corre E, Hourdez S, Bonnivard É, Claridge-Chang A, Jollivet D: Genomic patterns of divergence in the early and late steps of speciation of the deep-sea vent thermophilic worms of the genus Alvinella. BMC Ecol Evol 2022, 22:106.

25. Perez M, Aroh O, Sun Y, Lan Y, Juniper SK, Young CR, Angers B, Qian P-Y: Third-Generation Sequencing Reveals the Adaptive Role of the Epigenome in Three Deep-Sea Polychaetes. Mol Biol Evol 2023, 40.

26. Plouviez S, Shank TM, Faure B, Daguin-Thiebaut C, Viard F, Lallier FH, Jollivet D: Comparative phylogeography among hydrothermal vent species along the East Pacific Rise reveals vicariant processes and population expansion in the South. Mol Ecol 2009, 18:3903–3917.

27. Plouviez S, Le Guen D, Lecompte O, Lallier FH, Jollivet D: Determining gene flow and the influence of selection across the equatorial barrier of the East Pacific Rise in the tube-dwelling polychaete Alvinella pompejana. BMC Evol Biol 2010, 10:220.

28. Bonnivard E, Catrice O, Ravaux J, Brown SC, Higuet D: Survey of genome size in 28 hydrothermal vent species covering 10 families. Genome 2009, 52:524–536.

29. Dixon DR, Jolly MT, Vevers WF, Dixon LRJ: Chromosomes of Pacific hydrothermal vent invertebrates: towards a greater understanding of the relationship between chromosome and molecular evolution. J Mar Biol Assoc U K 2010, 90:15–31.

30. Martín-Durán JM, Vellutini BC, Marlétaz F, Cetrangolo V, Cvetesic N, Thiel D, Henriet S, Grau-Bové X, Carrillo-Baltodano AM, Gu W, et al.: Conservative route to genome compaction in a miniature annelid. Nat Ecol Evol 2021, 5:231–242.

31. Cary SC, Shank T, Stein J: Worms bask in extreme temperatures. Nature 1998, 391:545–546.

32. Dahlhoff E, O’Brien J, Somero GN, Vetter RD: Temperature Effects on Mitochondria from Hydrothermal Vent Invertebrates: Evidence for Adaptation to Elevated and Variable Habitat Temperatures. Physiol Zool 1991, 64:1490–1508.

33. Perez M, Wang H, Angers B, Qian P-Y: Complete mitochondrial genome of paralvinella palmiformis (Polychaeta: Alvinellidae). Mitochondrial DNA B Resour 2022, 7:786–788.

34. Weigert A, Golombek A, Gerth M, Schwarz F, Struck TH, Bleidorn C: Evolution of mitochondrial gene order in Annelida. Mol Phylogenet Evol 2016, 94:196–206.

35. Gu Z, Gu L, Eils R, Schlesner M, Brors B: circlize Implements and enhances circular visualization in R. Bioinformatics 2014, 30:2811–2812.

36. Lartillot N, Philippe H: A Bayesian mixture model for across-site heterogeneities in the amino-acid replacement process. Mol Biol Evol 2004, 21:1095–1109.

37. Struck TH, Paul C, Hill N, Hartmann S, Hösel C, Kube M, Lieb B, Meyer A, Tiedemann R, Purschke G, et al.: Phylogenomic analyses unravel annelid evolution. Nature 2011, 471:95–98.

38. Weigert A, Helm C, Meyer M, Nickel B, Arendt D, Hausdorf B, Santos SR, Halanych KM, Purschke G, Bleidorn C, et al.: Illuminating the base of the annelid tree using transcriptomics. Mol Biol Evol 2014, 31:1391–1401.

39. Marlétaz F, Peijnenburg KTCA, Goto T, Satoh N, Rokhsar DS: A New Spiralian Phylogeny Places the Enigmatic Arrow Worms among Gnathiferans. Curr Biol 2019, 29:312–318.e3.

40. Cottrell MT, Cary SC: Diversity of dissimilatory bisulfite reductase genes of bacteria associated with the deep-sea hydrothermal vent polychaete annelid Alvinella pompejana. Appl Environ Microbiol 1999, 65:1127–1132.

41. Grzymski JJ, Murray AE, Campbell BJ, Kaplarevic M, Gao GR, Lee C, Daniel R, Ghadiri A, Feldman RA, Cary SC: Metagenome analysis of an extreme microbial symbiosis reveals eurythermal adaptation and metabolic flexibility. Proc Natl Acad Sci U S A 2008, 105:17516–17521.

42. Payne SH, Loomis WF: Retention and loss of amino acid biosynthetic pathways based on analysis of whole-genome sequences. Eukaryot Cell 2006, 5:272–276.

43. Guedes RLM, Prosdocimi F, Fernandes GR, Moura LK, Ribeiro HAL, Ortega JM: Amino acids biosynthesis and nitrogen assimilation pathways: a great genomic deletion during eukaryotes evolution. BMC Genomics 2011, 12 Suppl 4:S2.

44. Richter DJ, Fozouni P, Eisen MB, King N: Gene family innovation, conservation and loss on the animal stem lineage. Elife 2018, 7.

45. Trolle J, McBee RM, Kaufman A, Pinglay S, Berger H, German S, Liu L, Shen MJ, Guo X, Martin JA, et al.: Resurrecting essential amino acid biosynthesis in mammalian cells. Elife 2022, 11.

46. Neave MJ, Michell CT, Apprill A, Voolstra CR: Endozoicomonas genomes reveal functional adaptation and plasticity in bacterial strains symbiotically associated with diverse marine hosts. Sci Rep 2017, 7:40579.

47. Ide K, Nishikawa Y, Maruyama T, Tsukada Y, Kogawa M, Takeda H, Ito H, Wagatsuma R, Miyaoka R, Nakano Y, et al.: Targeted single-cell genomics reveals novel host adaptation strategies of the symbiotic bacteria Endozoicomonas in Acropora tenuis coral. Microbiome 2022, 10:220.

48. Sueoka N: CORRELATION BETWEEN BASE COMPOSITION OF DEOXYRIBONUCLEIC ACID AND AMINO ACID COMPOSITION OF PROTEIN. Proc Natl Acad Sci U S A 1961, 47:1141–1149.

49. Kreil DP, Ouzounis CA: Identification of thermophilic species by the amino acid compositions deduced from their genomes. Nucleic Acids Res 2001, 29:1608–1615.

50. Simakov O, Bredeson J, Berkoff K, Marletaz F, Mitros T, Schultz DT, O’Connell BL, Dear P, Martinez DE, Steele RE, et al.: Deeply conserved synteny and the evolution of metazoan chromosomes. Sci Adv 2022, 8:eabi5884.

51. Wang S, Zhang J, Jiao W, Li J, Xun X, Sun Y, Guo X, Huan P, Dong B, Zhang L, et al.: Scallop genome provides insights into evolution of bilaterian karyotype and development. Nat Ecol Evol 2017, 1:120.

52. Hui JHL, McDougall C, Monteiro AS, Holland PWH, Arendt D, Balavoine G, Ferrier DEK: Extensive chordate and annelid macrosynteny reveals ancestral homeobox gene organization. Mol Biol Evol 2012, 29:157–165.

53. Simakov O, Kawashima T, Marlétaz F, Jenkins J, Koyanagi R, Mitros T, Hisata K, Bredeson J, Shoguchi E, Gyoja F, et al.: Hemichordate genomes and deuterostome origins. Nature 2015, 527:459–465.

54. Kapli P, Natsidis P, Leite DJ, Fursman M, Jeffrie N, Rahman IA, Philippe H, Copley RR, Telford MJ: Lack of support for Deuterostomia prompts reinterpretation of the first Bilateria. Sci Adv 2021, 7.

55. Wotton KR, Weierud FK, Juárez-Morales JL, Alvares LE, Dietrich S, Lewis KE: Conservation of gene linkage in dispersed vertebrate NK homeobox clusters. Dev Genes Evol 2009, 219:481–496.

56. Seudre O, Martín-Zamora FM, Rapisarda V, Luqman I, Carrillo-Baltodano AM, Martín-Durán JM: The Fox Gene Repertoire in the Annelid Owenia fusiformis Reveals Multiple Expansions of the foxQ2 Class in Spiralia. Genome Biol Evol 2022, 14.

57. Fritzenwanker JH, Gerhart J, Freeman RM Jr, Lowe CJ: The Fox/Forkhead transcription factor family of the hemichordate Saccoglossus kowalevskii. Evodevo 2014, 5:17.

58. Montoro DT, Haber AL, Biton M, Vinarsky V, Lin B, Birket SE, Yuan F, Chen S, Leung HM, Villoria J, et al.: A revised airway epithelial hierarchy includes CFTR-expressing ionocytes. Nature 2018, 560:319–324.

59. Sackville MA, Cameron CB, Gillis JA, Brauner CJ: Ion regulation at gills precedes gas exchange and the origin of vertebrates. Nature 2022, 610:699–703.

60. Gaill F, Bouligand Y: Supercoil of collagen fibrils in the integument of Alvinella, an abyssal annelid. Tissue Cell 1987, 19:625–642.

61. Taga Y, Tanaka K, Hattori S, Mizuno K: In-depth correlation analysis demonstrates that 4-hydroxyproline at the Yaa position of Gly-Xaa-Yaa repeats dominantly stabilizes collagen triple helix. Matrix Biology Plus 2021, 10:100067.

62. Sicot FX, Mesnage M, Masselot M, Exposito JY, Garrone R, Deutsch J, Gaill F: Molecular adaptation to an extreme environment: origin of the thermal stability of the pompeii worm collagen. J Mol Biol 2000, 302:811–820.

63. Taga Y, Kusubata M, Mizuno K: Quantitative Analysis of the Positional Distribution of Hydroxyproline in Collagenous Gly-Xaa-Yaa Sequences by LC-MS with Partial Acid Hydrolysis and Precolumn Derivatization. Anal Chem 2020, 92:8427–8434.

64. Gaill F, Hunt S: Tubes of deep sea hydrothermal vent worms Riftia pachyptila (Vestimentifera) and Alvinella pompejana (Annelida). Mar Ecol Prog Ser 1986, 34:267–274.

65. Vovelle J, Gaill F: Donńees morphologiques, histochimiques et microanalytiques sur l’élaboration du tube organominéral d’*Alvinella pompejana*, Polychète des sources hydrothermales, et leurs implications Phylogénétiques. Zool Scr 1986, 15:33–43.

66. Zhou CZ, Confalonieri F, Jacquet M, Perasso R, Li ZG, Janin J: Silk fibroin: structural implications of a remarkable amino acid sequence. Proteins 2001, 44:119–122.

67. Han Z, Wang Z, Rittschof D, Huang Z, Chen L, Hao H, Yao S, Su P, Huang M, Zhang Y-Y, et al.: New genes helped acorn barnacles adapt to a sessile lifestyle. Nat Genet 2024, 56:970–981.

68. Buffet J-P, Corre E, Duvernois-Berthet E, Fournier J, Lopez PJ: Adhesive gland transcriptomics uncovers a diversity of genes involved in glue formation in marine tube-building polychaetes. Acta Biomater 2018, 72:316–328.

69. Song S, Starunov V, Bailly X, Ruta C, Kerner P, Cornelissen AJM, Balavoine G: Globins in the marine annelid Platynereis dumerilii shed new light on hemoglobin evolution in bilaterians. BMC Evol Biol 2020, 20:165.

70. Belato FA, Coates CJ, Halanych KM, Weber RE, Costa-Paiva EM: Evolutionary History of the Globin Gene Family in Annelids. Genome Biol Evol 2020, 12:1719–1733.

71. Flores JF, Fisher CR, Carney SL, Green BN, Freytag JK, Schaeffer SW, Royer WE Jr: Sulfide binding is mediated by zinc ions discovered in the crystal structure of a hydrothermal vent tubeworm hemoglobin. Proc Natl Acad Sci U S A 2005, 102:2713–2718.

72. Kim J, Fukuda Y, Inoue T: Crystal structure of Kumaglobin: a hexacoordinated heme protein from an anhydrobiotic tardigrade, Ramazzottius varieornatus. FEBS J 2019, 286:1287–1304.

73. Bailly X, Leroy R, Carney S, Collin O, Zal F, Toulmond A, Jollivet D: The loss of the hemoglobin H_2_S-binding function in annelids from sulfide-free habitats reveals molecular adaptation driven by Darwinian positive selection. Proc Natl Acad Sci U S A 2003, 100:5885–5890.

74. Belato FA, Schrago CG, Coates CJ, Halanych KM, Costa-Paiva EM: Newly Discovered Occurrences and Gene Tree of the Extracellular Globins and Linker Chains from the Giant Hexagonal Bilayer Hemoglobin in Metazoans. Genome Biol Evol 2019, 11:597–612.

75. LaCount MW, Zhang E, Chen YP, Han K, Whitton MM, Lincoln DE, Woodin SA, Lebioda L: The Crystal Structure and Amino Acid Sequence of Dehaloperoxidase from Amphitrite ornata Indicate Common Ancestry with Globins*. J Biol Chem 2000, 275:18712–18716.

76. Bailly X, Chabasse C, Hourdez S, Dewilde S, Martial S, Moens L, Zal F: Globin gene family evolution and functional diversification in annelids. FEBS J 2007, 274:2641–2652.

77. Chen YP, Woodin SA, Lincoln DE, Lovell CR: An unusual dehalogenating peroxidase from the marine terebellid polychaete Amphitrite ornata. J Biol Chem 1996, 271:4609–4612.

78. Costa-Paiva EM, Whelan NV, Waits DS, Santos SR, Schrago CG, Halanych KM: Discovery and evolution of novel hemerythrin genes in annelid worms. BMC Evol Biol 2017, 17:85.

79. Hashimoto T, Horikawa DD, Saito Y, Kuwahara H, Kozuka-Hata H, Shin-I T, Minakuchi Y, Ohishi K, Motoyama A, Aizu T, et al.: Extremotolerant tardigrade genome and improved radiotolerance of human cultured cells by tardigrade-unique protein. Nat Commun 2016, 7:12808.

80. Graham AM, Barreto FS: Loss of the HIF pathway in a widely distributed intertidal crustacean, the copepod Tigriopus californicus. Proc Natl Acad Sci U S A 2019, 116:12913–12918.

81. Kaelin WG Jr, Ratcliffe PJ: Oxygen sensing by metazoans: the central role of the HIF hydroxylase pathway. Mol Cell 2008, 30:393–402.

82. Hug LA, Stechmann A, Roger AJ: Phylogenetic distributions and histories of proteins involved in anaerobic pyruvate metabolism in eukaryotes. Mol Biol Evol 2010, 27:311–324.

83. Kim S, Kim CM, Son Y-J, Choi JY, Siegenthaler RK, Lee Y, Jang T-H, Song J, Kang H, Kaiser CA, et al.: Molecular basis of maintaining an oxidizing environment under anaerobiosis by soluble fumarate reductase. Nat Commun 2018, 9:4867.

84. Müller M, Mentel M, van Hellemond JJ, Henze K, Woehle C, Gould SB, Yu R-Y, van der Giezen M, Tielens AGM, Martin WF: Biochemistry and evolution of anaerobic energy metabolism in eukaryotes. Microbiol Mol Biol Rev 2012, 76:444–495.

85. Tielens AGM, Rotte C, van Hellemond JJ, Martin W: Mitochondria as we don’t know them. Trends Biochem Sci 2002, 27:564–572.

86. Tan JH, Lautens M, Romanelli-Cedrez L, Wang J, Schertzberg MR, Reinl SR, Davis RE, Shepherd JN, Fraser AG, Salinas G: Alternative splicing of coq-2 controls the levels of rhodoquinone in animals. Elife 2020, 9.

87. Roberts Buceta PM, Romanelli-Cedrez L, Babcock SJ, Xun H, VonPaige ML, Higley TW, Schlatter TD, Davis DC, Drexelius JA, Culver JC, et al.: The kynurenine pathway is essential for rhodoquinone biosynthesis in Caenorhabditis elegans. J Biol Chem 2019, 294:11047–11053.

88. Del Borrello S, Lautens M, Dolan K, Tan JH, Davie T, Schertzberg MR, Spensley MA, Caudy AA, Fraser AG: Rhodoquinone biosynthesis in C. elegans requires precursors generated by the kynurenine pathway. Elife 2019, 8.

89. Moggioli G, Panossian B, Sun Y, Thiel D, Martín-Zamora FM, Tran M, Clifford AM, Goffredi SK, Rimskaya-Korsakova N, Jékelly G, et al.: The hologenome of Osedax frankpressi reveals the genetic interplay for the symbiotic digestion of vertebrate bone. bioRxiv 2022, doi:10.1101/2022.08.04.502725.

90. Sun J, Zhang Y, Xu T, Zhang Y, Mu H, Zhang Y, Lan Y, Fields CJ, Hui JHL, Zhang W, et al.: Adaptation to deep-sea chemosynthetic environments as revealed by mussel genomes. Nat Ecol Evol 2017, 1:121.

91. Mat AM, Sarrazin J, Markov GV, Apremont V, Dubreuil C, Eché C, Fabioux C, Klopp C, Sarradin P-M, Tanguy A, et al.: Biological rhythms in the deep-sea hydrothermal mussel Bathymodiolus azoricus. Nat Commun 2020, 11:3454.

92. Cuvelier D, Legendre P, Laes A, Sarradin P-M, Sarrazin J: Rhythms and community dynamics of a hydrothermal tubeworm assemblage at main endeavour field - a multidisciplinary deep-sea observatory approach. PLoS One 2014, 9:e96924.

93. Mi H, Ebert D, Muruganujan A, Mills C, Albou L-P, Mushayamaha T, Thomas PD: PANTHER version 16: a revised family classification, tree-based classification tool, enhancer regions and extensive API. Nucleic Acids Res 2021, 49:D394–D403.

94. Bauer S, Grossmann S, Vingron M, Robinson PN: Ontologizer 2.0--a multifunctional tool for GO term enrichment analysis and data exploration. Bioinformatics 2008, 24:1650–1651.

95. Zhu F-C, Sun J, Yan G-Y, Huang J-M, Chen C, He L-S: Insights into the strategy of micro-environmental adaptation: Transcriptomic analysis of two alvinocaridid shrimps at a hydrothermal vent. PLoS One 2020, 15:e0227587.

96. Dilly GF, Young CR, Lane WS, Pangilinan J, Girguis PR: Exploring the limit of metazoan thermal tolerance via comparative proteomics: thermally induced changes in protein abundance by two hydrothermal vent polychaetes. Proc Biol Sci 2012, 279:3347–3356.

97. Ip JC-H, Xu T, Sun J, Li R, Chen C, Lan Y, Han Z, Zhang H, Wei J, Wang H, et al.: Host-Endosymbiont Genome Integration in a Deep-Sea Chemosymbiotic Clam. Mol Biol Evol 2021, 38:502–518.

98. Sun J, Chen C, Miyamoto N, Li R, Sigwart JD, Xu T, Sun Y, Wong WC, Ip JCH, Zhang W, et al.: The Scaly-foot Snail genome and implications for the origins of biomineralised armour. Nat Commun 2020, 11:1657.

99. Lan Y, Sun J, Chen C, Sun Y, Zhou Y, Yang Y, Zhang W, Li R, Zhou K, Wong WC, et al.: Hologenome analysis reveals dual symbiosis in the deep-sea hydrothermal vent snail Gigantopelta aegis. Nat Commun 2021, 12:1165.

100. Zhou Y, Liu H, Feng C, Lu Z, Liu J, Huang Y, Tang H, Xu Z, Pu Y, Zhang H: Genetic adaptations of sea anemone to hydrothermal environment. Sci Adv 2023, 9:eadh0474.

101. Law STS, Yu Y, Nong W, So WL, Li Y, Swale T, Ferrier DEK, Qiu J, Qian P, Hui JHL: The genome of the deep-sea anemone Actinernus sp. contains a mega-array of ANTP-class homeobox genes. Proc Biol Sci 2023, 290:20231563.

102 Gagnaire P-A: Comparative genomics approach to evolutionary process connectivity. Evol Appl 2020, 13:1320–1334.

103. Castelin M, Lorion J, Brisset J, Cruaud C, Maestrati P, Utge J, Samadi S: Speciation patterns in gastropods with long-lived larvae from deep-sea seamounts. Mol Ecol 2012, 21:4828–4853.

104. Pante E, France SC, Couloux A, Cruaud C, McFadden CS, Samadi S, Watling L: Deep-sea origin and in-situ diversification of chrysogorgiid octocorals. PLoS One 2012, 7:e38357.

105. Castel J, Hourdez S, Pradillon F, Daguin-Thiébaut C, Ballenghien M, Ruault S, Corre E, Tran Lu Y A, Mary J, Gagnaire P-A, et al.: Inter-Specific Genetic Exchange Despite Strong Divergence in Deep-Sea Hydrothermal Vent Gastropods of the Genus Alviniconcha. Genes 2022, 13.

106. Wu C: The genic view of the process of speciation. J Evol Biol 2001, 14:851–865.

107. Le Moan A, Roby C, Fraïsse C, Daguin-Thiébaut C, Bierne N, Viard F: An introgression breakthrough left by an anthropogenic contact between two ascidians. Mol Ecol 2021, 30:6718–6732.

108. Fraïsse C, Le Moan A, Roux C, Dubois G, Daguin-Thiebaut C, Gagnaire P-A, Viard F, Bierne N: Introgression between highly divergent sea squirt genomes: an adaptive breakthrough? Peer Community J 2022, 2.

109. Plouviez S, Faure B, Le Guen D, Lallier FH, Bierne N, Jollivet D: A new barrier to dispersal trapped old genetic clines that escaped the Easter Microplate tension zone of the Pacific vent mussels. PLoS One 2013, 8:e81555.

110. Jarzab A, Kurzawa N, Hopf T, Moerch M, Zecha J, Leijten N, Bian Y, Musiol E, Maschberger M, Stoehr G, et al.: Meltome atlas-thermal proteome stability across the tree of life. Nat Methods 2020, 17:495–503.

111. Le Bris N, Gaill F: How does the annelid Alvinella pompejana deal with an extreme hydrothermal environment? Rev Environ Sci Technol 2007, 6:197–221.

112. Mentel M, Martin W: Anaerobic animals from an ancient, anoxic ecological niche. BMC Biol 2010, 8:32.

113. Ruan J, Li H: Fast and accurate long-read assembly with wtdbg2. Nat Methods 2020, 17:155–158.

114. Putnam NH, O’Connell BL, Stites JC, Rice BJ, Blanchette M, Calef R, Troll CJ, Fields A, Hartley PD, Sugnet CW, et al.: Chromosome-scale shotgun assembly using an in vitro method for long-range linkage. Genome Res 2016, 26:342–350.

115. Lieberman-Aiden E, van Berkum NL, Williams L, Imakaev M, Ragoczy T, Telling A, Amit I, Lajoie BR, Sabo PJ, Dorschner MO, et al.: Comprehensive mapping of long-range interactions reveals folding principles of the human genome. Science 2009, 326:289–293.

116. Flynn JM, Hubley R, Goubert C, Rosen J, Clark AG, Feschotte C, Smit AF: RepeatModeler2 for automated genomic discovery of transposable element families. Proc Natl Acad Sci U S A 2020, 117:9451–9457.

117. Dobin A, Davis CA, Schlesinger F, Drenkow J, Zaleski C, Jha S, Batut P, Chaisson M, Gingeras TR: STAR: ultrafast universal RNA-seq aligner. Bioinformatics 2013, 29:15–21.

118. Kovaka S, Zimin AV, Pertea GM, Razaghi R, Salzberg SL, Pertea M: Transcriptome assembly from long-read RNA-seq alignments with StringTie2. Genome Biol 2019, 20:278.

119. Grabherr MG, Haas BJ, Yassour M, Levin JZ, Thompson DA, Amit I, Adiconis X, Fan L, Raychowdhury R, Zeng Q, et al.: Full-length transcriptome assembly from RNA-Seq data without a reference genome. Nat Biotechnol 2011, 29:644–652.

120. Pertea G, Huang X, Liang F, Antonescu V, Sultana R, Karamycheva S, Lee Y, White J, Cheung F, Parvizi B, et al.: TIGR Gene Indices clustering tools (TGICL): a software system for fast clustering of large EST datasets. Bioinformatics 2003, 19:651–652.

121. Haas BJ, Delcher AL, Mount SM, Wortman JR, Smith RK Jr, Hannick LI, Maiti R, Ronning CM, Rusch DB, Town CD, et al.: Improving the Arabidopsis genome annotation using maximal transcript alignment assemblies. Nucleic Acids Res 2003, 31:5654–5666.

122. Keller O, Kollmar M, Stanke M, Waack S: A novel hybrid gene prediction method employing protein multiple sequence alignments. Bioinformatics 2011, 27:757–763.

123. Hoff KJ, Stanke M: Predicting Genes in Single Genomes with AUGUSTUS. Curr Protoc Bioinformatics 2019, 65:e57.

124. Haas BJ, Salzberg SL, Zhu W, Pertea M, Allen JE, Orvis J, White O, Buell CR, Wortman JR: Automated eukaryotic gene structure annotation using EVidenceModeler and the Program to Assemble Spliced Alignments. Genome Biol 2008, 9:R7.

125. Eddy SR: Accelerated Profile HMM Searches. PLoS Comput Biol 2011, 7:e1002195.

126. Mistry J, Chuguransky S, Williams L, Qureshi M, Salazar GA, Sonnhammer ELL, Tosatto SCE, Paladin L, Raj S, Richardson LJ, et al.: Pfam: The protein families database in 2021. Nucleic Acids Res 2021, 49:D412–D419.

127. Almagro Armenteros JJ, Tsirigos KD, Sønderby CK, Petersen TN, Winther O, Brunak S, von Heijne G, Nielsen H: SignalP 5.0 improves signal peptide predictions using deep neural networks. Nat Biotechnol 2019, 37:420–423.

128. Buchfink B, Xie C, Huson DH: Fast and sensitive protein alignment using DIAMOND. Nat Methods 2015, 12:59–60.

129. O’Leary NA, Wright MW, Brister JR, Ciufo S, Haddad D, McVeigh R, Rajput B, Robbertse B, Smith-White B, Ako-Adjei D, et al.: Reference sequence (RefSeq) database at NCBI: current status, taxonomic expansion, and functional annotation. Nucleic Acids Res 2016, 44:D733–45.

130. Kanehisa M, Sato Y, Morishima K: BlastKOALA and GhostKOALA: KEGG Tools for Functional Characterization of Genome and Metagenome Sequences. J Mol Biol 2016, 428:726–731.

131. Jin J-J, Yu W-B, Yang J-B, Song Y, dePamphilis CW, Yi T-S, Li D-Z: GetOrganelle: a fast and versatile toolkit for accurate de novo assembly of organelle genomes. Genome Biol 2020, 21:241.

132. Langmead B, Salzberg SL: Fast gapped-read alignment with Bowtie 2. Nat Methods 2012, 9:357–359.

133. Bankevich A, Nurk S, Antipov D, Gurevich AA, Dvorkin M, Kulikov AS, Lesin VM, Nikolenko SI, Pham S, Prjibelski AD, et al.: SPAdes: a new genome assembly algorithm and its applications to single-cell sequencing. J Comput Biol 2012, 19:455–477.

134. Camacho C, Coulouris G, Avagyan V, Ma N, Papadopoulos J, Bealer K, Madden TL: BLAST+: architecture and applications. BMC Bioinformatics 2009, 10:421.

135. Wick RR, Schultz MB, Zobel J, Holt KE: Bandage: interactive visualization of de novo genome assemblies. Bioinformatics 2015, 31:3350–3352.

136. Bernt M, Donath A, Jühling F, Externbrink F, Florentz C, Fritzsch G, Pütz J, Middendorf M, Stadler PF: MITOS: improved de novo metazoan mitochondrial genome annotation. Mol Phylogenet Evol 2013, 69:313–319.

137. Doyle JJ, Dickson EE: Preservation of plant samples for DNA restriction endonuclease analysis. Taxon 1987, 36:715–722.

138. Jolly M, Viard F, Weinmayr G, Gentil F: Does the genetic structure of Pectinaria koreni (Polychaeta: Pectinariidae) conform to a source–sink metapopulation model at the scale of the Baie de Seine? Helgol Mar Res 2003,

139. Reeve J, Butlin RK, Koch EL, Stankowski S, Faria R: Chromosomal inversion polymorphisms are widespread across the species ranges of rough periwinkles (Littorina saxatilis and L. arcana). Mol Ecol 2023, doi:10.1111/mec.17160.

140. Li H, Durbin R: Fast and accurate short read alignment with Burrows–Wheeler transform. Bioinformatics 2009, 25:1754–1760.

141. Danecek P, Bonfield JK, Liddle J, Marshall J, Ohan V, Pollard MO, Whitwham A, Keane T, McCarthy SA, Davies RM, et al.: Twelve years of SAMtools and BCFtools. Gigascience 2021, 10.

142. Danecek P, Auton A, Abecasis G, Albers CA, Banks E, DePristo MA, Handsaker RE, Lunter G, Marth GT, Sherry ST, et al.: The variant call format and VCFtools. Bioinformatics 2011, 27:2156–2158.

143. Jombart T: adegenet: a R package for the multivariate analysis of genetic markers. Bioinformatics 2008, 24:1403–1405.

144. Schliep KP: phangorn: phylogenetic analysis in R. Bioinformatics 2011, 27:592–593.

145. Simakov O, Marletaz F, Cho S-J, Edsinger-Gonzales E, Havlak P, Hellsten U, Kuo D-H, Larsson T, Lv J, Arendt D, et al.: Insights into bilaterian evolution from three spiralian genomes. Nature 2013, 493:526–531.

146. Pearson WR, Lipman DJ: Improved tools for biological sequence comparison. Proc Natl Acad Sci U S A 1988, 85:2444–2448.

147. Altenhoff AM, Levy J, Zarowiecki M, Tomiczek B, Warwick Vesztrocy A, Dalquen DA, Müller S, Telford MJ, Glover NM, Dylus D, et al.: OMA standalone: orthology inference among public and custom genomes and transcriptomes. Genome Res 2019, 29:1152–1163.

148. Katoh K, Misawa K, Kuma K-I, Miyata T: MAFFT: a novel method for rapid multiple sequence alignment based on fast Fourier transform. Nucleic Acids Res 2002, 30:3059–3066.

149. Capella-Gutiérrez S, Silla-Martínez JM, Gabaldón T: trimAl: a tool for automated alignment trimming in large-scale phylogenetic analyses. Bioinformatics 2009, 25:1972–1973.

150. Lartillot N, Lepage T, Blanquart S: PhyloBayes 3: a Bayesian software package for phylogenetic reconstruction and molecular dating. Bioinformatics 2009, 25:2286–2288.

151. El Hilali S: SamiLhll/rbhXpress: v1.2.3. 2022.

152. El Hilali S, Copley RR: macrosyntR : Drawing automatically ordered Oxford Grids from standard genomic files in R. bioRxiv 2023, doi:10.1101/2023.01.26.525673.

153. Emms DM, Kelly S: OrthoFinder: phylogenetic orthology inference for comparative genomics. Genome Biol 2019, 20:238.

154. Li H, Handsaker B, Wysoker A, Fennell T, Ruan J, Homer N, Marth G, Abecasis G, Durbin R, 1000 Genome Project Data Processing Subgroup: The Sequence Alignment/Map format and SAMtools. Bioinformatics 2009, 25:2078–2079.

155. Martin M: Cutadapt removes adapter sequences from high-throughput sequencing reads. EMBnet.journal 2011, 17:10–12.

156. Nurk S, Meleshko D, Korobeynikov A, Pevzner PA: metaSPAdes: a new versatile metagenomic assembler. Genome Res 2017, 27:824–834.

157. Gurevich A, Saveliev V, Vyahhi N, Tesler G: QUAST: quality assessment tool for genome assemblies. Bioinformatics 2013, 29:1072–1075.

158. Kang DD, Li F, Kirton E, Thomas A, Egan R, An H, Wang Z: MetaBAT 2: an adaptive binning algorithm for robust and efficient genome reconstruction from metagenome assemblies. PeerJ 2019, 7:e7359.

159. Parks DH, Imelfort M, Skennerton CT, Hugenholtz P, Tyson GW: CheckM: assessing the quality of microbial genomes recovered from isolates, single cells, and metagenomes. Genome Res 2015, 25:1043–1055.

160. Nguyen L-T, Schmidt HA, von Haeseler A, Minh BQ: IQ-TREE: a fast and effective stochastic algorithm for estimating maximum-likelihood phylogenies. Mol Biol Evol 2015, 32:268–274.

