## supplementary information including extra figures and some sequences for "Chromosome-scale genome assembly and gene annotation of the hydrothermal vent annelid *Alvinella pompejana* yield insight into animal evolution in extreme environments"

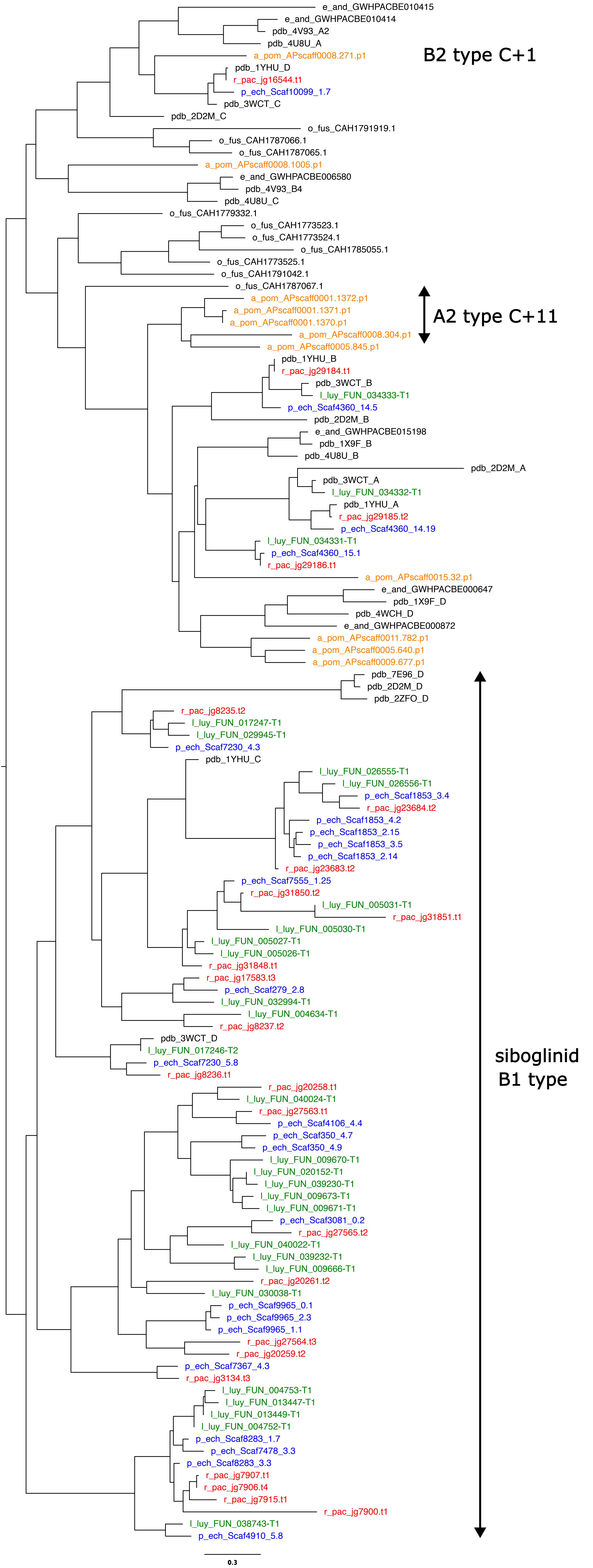
e_and_GWHPACBE010415
e_and_GWHPACBE010414
pdb_4V93_A2
pdb_4U8U_A
a_pom_APscaff0008.271.p1
pdb_1YHU_D
r_pac_jg16544.t1
p_ech_Scaf10099_1.7
pdb_3WCT_C
pdb_2D2M_C
B2 type C+1
o_fus_CAH1791919.1
o_fus_CAH1787066.1
o_fus_CAH1787065.1
a_pom_APscaff0008.1005.p1
e_and_GWHPACBE006580
pdb_4V93_B4
pdb_4U8U_C
o_fus_CAH1779332.1
o_fus_CAH1773523.1
o_fus_CAH1773524.1
o_fus_CAH1785055.1
o_fus_CAH1773525.1
o_fus_CAH1791042.1
o_fus_CAH1787067.1
a_pom_APscaff0001.1372.p1
a_pom_APscaff0001.1371.p1
a_pom_APscaff0001.1370.p1
a_pom_APscaff0008.304.p1
a_pom_APscaff0005.845.p1
A2 type C+11
pdb_1YHU_B
r_pac_jg29184.t1
pdb_3WCT_B
l_luy_FUN_034333-T1
p_ech_Scaf4360_14.5
pdb_2D2M_B
e_and_GWHPACBE015198
pdb_1X9F_B
pdb_4U8U_B
pdb_2D2M_A
pdb_3WCT_A
l_luy_FUN_034332-T1
pdb_1YHU_A
r_pac_jg29185.t2
p_ech_Scaf4360_14.19
l_luy_FUN_034331-T1
p_ech_Scaf4360_15.1
r_pac_jg29186.t1
a_pom_APscaff0015.32.p1
e_and_GWHPACBE000647
pdb_1X9F_D
pdb_4WCH_D
e_and_GWHPACBE000872
a_pom_APscaff0011.782.p1
a_pom_APscaff0005.640.p1
a_pom_APscaff0009.677.p1
pdb_7E96_D
pdb_2D2M_D
pdb_2ZFO_D
r_pac_jg8235.t2
l_luy_FUN_017247-T1
l_luy_FUN_029945-T1
p_ech_Scaf7230_4.3
pdb_1YHU_C
l_luy_FUN_026555-T1
l_luy_FUN_026556-T1
p_ech_Scaf1853_3.4
r_pac_jg23684.t2
p_ech_Scaf1853_4.2
p_ech_Scaf1853_2.15
p_ech_Scaf1853_3.5
p_ech_Scaf1853_2.14
r_pac_jg23683.t2
p_ech_Scaf7555_1.25
r_pac_jg31850.t2
l_luy_FUN_005031-T1
r_pac_jg31851.t1
l_luy_FUN_005030-T1
l_luy_FUN_005027-T1
l_luy_FUN_005026-T1
r_pac_jg31848.t1
r_pac_jg17583.t3
p_ech_Scaf279_2.8
l_luy_FUN_032994-T1
l_luy_FUN_004634-T1
r_pac_jg8237.t2
pdb_3WCT_D
l_luy_FUN_017246-T2
p_ech_Scaf7230_5.8
r_pac_jg8236.t1
siboglinid B1 type
r_pac_jg20258.t1
l_luy_FUN_040024-T1
r_pac_jg27563.t1
p_ech_Scaf4106_4.4
p_ech_Scaf350_4.7
p_ech_Scaf350_4.9
l_luy_FUN_009670-T1
l_luy_FUN_020152-T1
l_luy_FUN_039230-T1
l_luy_FUN_009673-T1
l_luy_FUN_009671-T1
p_ech_Scaf3081_0.2
r_pac_jg27565.t2
l_luy_FUN_040022-T1
l_luy_FUN_039232-T1
l_luy_FUN_009666-T1
r_pac_jg20261.t2
l_luy_FUN_030038-T1
p_ech_Scaf9965_0.1
p_ech_Scaf9965_2.3
p_ech_Scaf9965_1.1
r_pac_jg27564.t3
r_pac_jg20259.t2
p_ech_Scaf7367_4.3
r_pac_jg3134.t3
l_luy_FUN_004753-T1
l_luy_FUN_013447-T1
l_luy_FUN_013449-T1
l_luy_FUN_004752-T1
p_ech_Scaf8283_1.7
p_ech_Scaf7478_3.3
p_ech_Scaf8283_3.3
r_pac_jg7907.t1
r_pac_jg7906.t4
r_pac_jg7915.t1
r_pac_jg7900.t1
l_luy_FUN_038743-T1
p_ech_Scaf4910_5.8
0.3

*Miscellaneous protein sequences for genes mentioned in text.*

*Alvinella* **Antp** HOX:

```
>a_pom|APscaff0012.403.p1
MSSFYNSPYFDSELNNNFASYSALHYHGLARQNLALQQEFGDPYGNGVDQAAVPHPGTSY
PRFPPYERLDQIRPITSHGGGGGGGGGGGGGGGGGASTHNHMYQQGGHFGLTSPADMGHV
TSAIPNSTKQVASPPPPPPPRAHSSTPARQASPLGRVHGRSDVSPVLPDPIRETGSRDQI
HDHPHPTSVGLYCDNAVKMSVHDNKTDSRATIGTRSPEGSLPAAKDPPANHSPACAEKEK
QSAAGAGAVKEAPSGGESQLPPAESKKSLTDIPDFLSNCKIKDEDIEKMSSMFNDPEMSP
INHRADTPSAAHAQPGTPKSQSAAEDEPGQGQAQTGQGQKNGDSSDEKANSDDGKLRSPE
GAEGDSKSDDNMDEKKDSNSIPMYPWMRSQFGPERKRGRQTYTRYQTLELEKEFHFNRYL
TRRRRIEIAHALCLTERQIKIWFQNRRMKWKKETKQLELLRQAGELPDGLFDDK
```

*Alvinella* pyruvate:NADP+ oxidoreductase (PNO):

```
>a_pom|APscaff0002.389.p1  GENE.evm.model.APscaff0002.389~~evm.model.APscaff0002.389.p
1  ORF type:complete len:1822 (+),score=442.00 evm.model.APscaff0002.389:1-5466(+)
MWRAWIVGRPVLQTVMRAGCSSRRGLAEITTPRPAMAAVAPAQVKGRPEIRRNENQPWVV
MDGNEAAAYVAYQMSDISFIYPISPATSMGEHMDKWAAQGRKNILGQVVDVNMMQSEAGA
AGALHGAAAAGTLTSTFTASQGLLLMIPNMYLLAGELMPTVFHVSARTVSKHALSIFNDH
SDVMATRQTGFSMLCSASVQEVMDLGVAAHISALKSRLPFLHFFDGYRTSAEMSKIRMMP
PEDIQQIFPYEQVKEHLQKYALNPNSPSIRGTGQRPDIFFQTTVAANRFYNQCPDVVEET
FDEISALTGRKYGLFSYHGSPEADRVAVCMGSASKTLQETVDYLNERGEKTGVVTVHLFR
PWSTKHFLEVLPPSVSKIAVLDRTREDGAVGMPLFLDVNVTMSDAGRNTLITGGQYGLAS
KEFTPAMAKGVFDNLNAPQPKMRYVIGIEDDVTHTHLPYGENIRTTPKSIRQCLFWGLGS
DGTVGANKTAIKTIGLNTDMNAQGHFVYDSHKGDGVTVSHLRFGPEEIKSEYTIQNDADY
LSCSHPSYVYRYEMLEPLKEGGTFVLNSPWTTLEQMEKKLPAHIKNEIAQKKLQFYNIDA
TAIAQSVGLGKRVNMIMQAAFYGLAGVLPQDEAVNLLKKSIETQYSHKGPKVIEMNHKAV
DATMENLTKIQYPESWKDTTEGGSRPGVGNVKKPEFVTNIMDPVLALEGDKLPVSAFVPG
GYQPTGTTKYEKRGIAPAIPVWKPDACTQCNYCSIVCPHAVIRPFLLNKEENKKIPPGFE
ARKAKGGAEVAGYHYTIQVSPYDCTGCEVCVQSCPDDALYMAPFNEVADTYAPHWDYAIS
LPEREEVGDKYTVKGSQFMKPLFEFSGACAGCGETPYLKLATQLFGERMVIANASGCSSV
WGGTSTTIPFSTNREGRGPAWGRSLFEDNAEYGFGMMLATKQRRLKLRQEISAALLEMEL
SDKMKSLFNSYLQYFENPDKCDEVSAAIIAEFEAIGKDNLDPKLRSIYEQRDMLRTQSHW
LVGGDGWAYDIGYGGLDHVFSRGENVNILVLDTEMYSNTGGQVSKATQLSTVAKFATKGK
RQVKKDLGLCAMQYENVYVASVALGANMNQCVQAFKEAEHYNGTSLIIAYAPCIDWGIEM
KNMMKEMKRAVDTGYWSLYRYDPRRAEKGLNPFQLDSKKIKADLEAYLDGQNRFQQLKRS
DKDVAGQLHQHLSADIHHKHDKLMKMSMDNYELLEHLKNLLGETTDTEKVVVLYGSETGN
SAALADVFANELKRRGLRPKCMAMDDFDFDDLPKQDKVFCVVATCGQGEFPGNCKEFWKQ
VSDKELPKDFLKDVQVAVFGMCDRSYVYYNSAAKAFKERFEQLGAKSVMPLGYGDEKDED
RYETAWNEWLPELWNELGTGPPSQELLPPTYSVTMDATGLTVPDVIVPRGSKLLPMMKNV
VLTPPDYDRDIRHYEFDLSGSGFSYSVGDCLGIYPHNNKEDVLKFLDDYGLHSDMVISVQ
DTQGRKDPLPENTTISQLFTEVLDIFGKPARRFYETLSIAAKDEKEKSELEFLLSKDGKD
KLKELTKETVTYADLLRMYPSTKLSLEYLLDHVSRIRPRLYSIASSSEMFGDMLHLCIVK
DDWVTPSGKYRQGLCTRYLQGLSQGSTPDLVAGKMNAAGINIPDSQAPPYVMVALGTGIA
PMRAMIQDREVARMRGESVGPMALFFGARHKRTDYTYGDEFEEWHSGGKGVLNVLSTAFS
RDQAHKIYVQHRIAEHPELIYDYLWKRKGYFYLCGPAGNVPMSVRKAVVDAFVSQGGHSL
AEADKMVTQMQIEGRYNVEAW*
```

*Alvinella* fumarate reductase (*sensu* yeast OSM1 / FRD):

```
>A_pom|Gene.16723::CL5714Contig1::g.16723::m.16723
Gene.16723::CL5714Contig1::g.16723  ORF type:complete len:516 (+) CL5714Contig1:290-
1837(+)
MSTSSSQTTERVIVVGGGLAGLSAAVEASRHGAKVTIVEKEKQLGGNSAKATSGINGVGT
EAQSAKAIVDDVARFVEDTTKSGAGQSKQELVQVLGRNSAEAHVWLKSFGLNLTDVVQLG
GHSVPRTHRFPPTPDGKPIPVGFTIVSTLRKEVETKLKETVTIVTNAVFKKLLTDGDAVV
GVQYSDESGKLHDVEGTVVLAAGGYANDHTSDSLLVKHVPDLAKLPTTNGPWATGDIIKA
TADLSLSLVNMDRVQVHPTGFIEPKAPNEHTKFLAPEALRGCGAILLDSSGKRFVNELGR
RNYVSDSIFKHGKPYQGNDEYPVVAAMLLTQAVIDKFGPPAIGFYKFKGLIEDVNNLDGV
```

AQKMGVDVAVLKDTIKQYEADAKTGKDQFGKEDFPTVFSENDHFFLAYVTPTLHYCMGGI
EINTDANVLRPGSRIVPGLYAAGEVSGGVHGVNRLGGNSLLECVVFGRIAGRNAAHK*

Pfam **Octopine_DH** containing *Alvinella* sequences:

>a_pom|APscaff0002.759.p1 GENE.evm.model.APscaff0002.759~~evm.model.APscaff0002.759.p1 ORF type:complete len:405 (+),score=35.73 evm.model.APscaff0002.759:1-1215(+)
MITVLVCGGGNGAHVCAGLGASHDNVTTRVLTLYEDEAEKWTKAMGQDGIRITLRHSESD
CLTVVGSPALVTKRAEEAMKPNVDLIIITVPSFAHEQYLKALKPYVKPGMVIVGCPGRAG
FDFAVRSIWAELWSQVSIMNMESLPWACRISKFGCSVDVLGVKETLAGAVQKGEAPTRSA
LDPADMFQKVLGERPRLLTRGHLLGVTLSPTNGCIHPEIMYGRWKDWDGQPMNEPPLFYN
GLDRDTAELISAVSDEVMEIARAIMRQRPQVDLTNVEHIYQWYLRTYPDDIQDKSTLYTS
IRTNKAYKGLVHPCKETEDGRYVPNFKHRYLTEDLPYGMIVLKGIAEVAGVDTPRMDQVI
VWAQRKIGRSFIVGGSLTGEDLDITRSPQRYDFNTLDAITGLTN*
>a_pom|APscaff0002.1541.p1 GENE.evm.model.APscaff0002.1541~~evm.model.APscaff0002.1541.p1 ORF type:complete len:402 (+),score=73.63 evm.model.APscaff0002.1541:1-1206(+)
MVVAVICGGGNGAHVCAGIAASQPGVEARVLTLFADEAERWTNSLKEHDFTVTVHAAKKE
PTKLVAKPTMVTKVPGDAMQGSVDIILFTVPAFAHKQYLEELKPHVKPGMILAGCPGQAG
FEFAVRGIWGDLALQVSVLSFESLPWACRILEFGKSAEVLGTKGTLVGAVSESNPPPKSD
PTATLQKVLGDAPKLVAKGHLLGITLMGTNGYLHPSIMYGKWHKWDGKPFNEVPIFYNGL
DEFSAQVLSDISDEVVATAKAIMEQRPKVDLNNVSHILQWYHRCYGEDIEDKSTLYTCIR
TNRAYKGLTHPCVKNDDGTYVPNFKYRYLTEDIPFGLVVMRGIASIAGVQTPNMDKVITW
AQKQLGKEYLVDGQLKGNDLNETRCPQRYGLESLDKVLGLA*

*Alvinella* **urea cycle** proteins:

*argininosuccinate lyase:*

>a_pom|APscaff0004.737.p1 (**ASL1**)
GENE.evm.model.APscaff0004.737~~evm.model.APscaff0004.737.p1 ORF type:complete
len:578 (+),score=116.81 evm.model.APscaff0004.737:1-1734(+)
MRRRRLNMSMWKECSTLSATTSEKSATSNNRRKRRNFFDEIQQLMPSCVKRLHFNNEQEV
PRYAECLDLSEESIDQSDEDGDAGDSGGGGGGEQLEECFSCKVLDAINKASTKMAETNKG
GKLWGGRFTGTTDPIMHEFNASISYDKCMWKADIQGSKAWVSALLKAGLVTEEEKELITS
GLSKISDEWAAGTFSLEPTDEDIHTANERRLKELIGPVGGKLHTGRSRNDQVSTDMRLWL
RESIGNMKNLLKTLIAVFVSRAKREISILMPGYTHLQRAQPIRWSHWLLSYASMLQRDYE
RLDSLTPRVNTLTLGSGALAGNPFNIDMNLLAEKLGMERISLNSLDAASDRDFIAEFLFW
ASLTSVHLSRWAEDLILYSTAEFGFVTMSDAYSTGSSMMPQKKNADSLELIRGKAGHVYG
QCTILMVTMKGLPSTYNKDLQEDKQAMFDVYDTLTGVMQVAAGVLSTLKINADKCQRALS
LDMLATDIAYYLVRKGMPFREAHSLSGKCVALAEKRGCTLDKLSLKEFNDIHPLFTEDVM
KLWDFESSVEQYQSPNGTSSSSVLSQINILETWLNSK*

*argininosuccinate synthetase 1:*

>a_pom|APscaff0003.931.p1
(**ASS1**) GENE.evm.model.APscaff0003.931~~evm.model.APscaff0003.931.p1 ORF
type:complete len:410 (+),score=67.30 evm.model.APscaff0003.931:1-1230(+)
MSTKDTVVLAYSGGLDTSCILKWLQEKGYDVITFTANVGQDEDFDAARAKAEKLGAKKVV
IQDLRQEFFEEFISVGIQANAIYEDRYLMGTAFARPCIARAIVKVAKEEGAKYISHGATG
KGNDQVRFELACYALYPEVKLISPWRLPEFYTRFRGRPDLFKYAEEHGIPLPVSPKAPWS
IDANMMHVSYESGILEDPRNEAPATLYEMTTDPTKAPDSPERLVIEFKNGIPVKVKNLND
NTEISGGLNLYMYLNKIGSRHGVGRIDIVENRFLGMKSRGIYETPAGTILYQAHLDIENL
TMDRELRKIKQQLSVQFSEQVYRGFWFSPECAFVRHCIAKSQEGVDGTVYVKVYKGNVYI
TSRESACSLYNQELVSMDVQGNYDPSDAAGFIIVNALRLKEYNRRQIQK*

*arginase:*

>a_pom|APscaff0004.221.p1 (**ARG1/2**)
GENE.evm.model.APscaff0004.221~~evm.model.APscaff0004.221.p1 ORF type:complete
len:322 (+),score=25.06 evm.model.APscaff0004.221:1-966(+)

MSVFDKPVGVVGIPFDKGQPRKGVNHGPEVLRRAGAVSIIEELGYDVTDYGDIRLEDVPN
DPPAFGVKLPRTIGGDMKRLSDKVSQVVASGAICLNLGGDHTLGIGSINGHLAAKPHAAV
IWIDAHADLNPPTSSPSGNIHGMPLAFLIKEVQRYMPQLPGFEWLKARLNAQDVAYIGIR
DVDKAEKKLIKELGITYYDMYDIDRMGIHQVVEGALNAVNPRNDRPIHLSFDIDALDACY
CPSTGTPVSAGLTLREGMYIVEKIFRTGNLSVFDIAEVNPKLGSPEDVAVTVKSALSLIG
AAFGQRTVDFLADDYTIPKPN*

*Cluster of intracellular globins:*

>a_pom|APscaff0005.650.p1
MGCAPSKEGATFTSSDPQPANSSQSRSSGASKSILREAVESGSYSKMASYKPDPRCPLTE
RQLYSITKSWKAINREMASTAVNMFVRLLEFDGIRSFFSKFKDTQTVAELRANKVFEGHA
LSVISIIDEVITNLDDMDYVISLLQATGESHSIKFENFNPDFLWKVEGAFLWAVKETLGD
RYTISIENIYTTTIRYIIQSLHDAFTKHRKENPRESTETEKAGELPTNETETHGETKAIS
DNSLSVDFIGTNGATKHFVAAEE*
>a_pom|APscaff0005.649.p1
MGCAPSKEGATVKARDVHPATNSLSRSTDASKSILREAVETGSYSKMASYKPDPRCPLTE
RQLYSITKSWKAINREMASTALNMFVRLLEVRGIRAVFSKFKDHTTVAELRADAIFQSHA
LSVISVIDEVITNLDDMDYVISLLQATGESHSIQTENFQADYLWNVEGAFLWAVKETLGD
RYTISIEQIYVVTIRYILKSLHASFTEHRANRARAPTGGKNTQ*
>a_pom|APscaff0005.651.p1
MGCAPSKEGATVKAVDLQAVNSSQSRSSGASKSILREAVETGSYSKMASYKPDPRCPLTE
RQLYSITKSWKAINREMASTAVNMFIRLLEHDGIRSFFTKFKDHKTVAELRASKVFESHA
LMVISVIDDVITNLDDMDYVMSLLQATGESHSIKFKNFNPDFLWNVEGAFLWAVKETLGD
RYTISIENIYTITIRYILQSLHDAFTKHRERQNSTNNDSKKTNLLNQELSTADRKTAPDS
KD*
>a_pom|APscaff0005.652.p1
MGCAPSKEGATVTSGDTQRANGSQSRSSCATKSILREAVESGSYSKMASYKPDPRCPLTE
RQLYSITKSWKAINREMASTAVNMFVRLLEVDGMRSFFSKFKDHKTVAELRASKVFETHA
LMVISVIDDVITNLDDMDYVISLLEATGESHSKKFGNLNADFLWNVEEPFLWAIRETLGD
RYTISIENIYTITIRYILQALHDSFTKHRQGQTTAETETRSNEKQAEVEDTIGEKDKDLP
YDRMSSTEIKGKYYDEEIKPKLSVTSDKPKETVRN*
>a_pom|APscaff0005.653.p1
MGCAPSIEGATVTSGDVQPAKSSLSKSTGATKSILREAVDSGRYSRMASYKPDPRCPLTE
RQLYSITKSWKAINREMASTAVNMFVRLLEIDGIRSFFSKFKDHKTVAELRASEVFEGHA
LMVISAIDDVITNLDDMDYVISLLEETGQSHSRRFHSFNPAFFWKVEGAFLWAVKETLGD
RYTISIENIYTTTIRYILQSLHDAFVGHRGNHADDDPAKSESLLQHQEKKDEEEAAGPEK
PRTNEE*

*Protein with S/G/A amino acid composition percentages most similar to tube:*

>a_pom|APscaff0012.608.p1
MSVSAGCGKNHFSKSTSASGSYAIAQGKNGVCTVTASFSQSTSASAGCGKNHWSKSGSAS
GSYAMKKGKGGKLICKTGSYSCSTSVSAGVGKNHFSKSTSASGSYAVTQGKNGVCTCIAS
FSRSVSTSAGCGKNHYSMSASASGSVAKKTGKGGKLICQTGRFSSSKSVSAGVGKNHFSK
STSASGSYAIQKGKNGVCTVVSSFSRSVSTSVGCGKNHYSMSGSASGSFAKKTGKGGKLI
CQTGRYSSSTSVSAGVGKNHYSKSASLSTSYAIQKGKNGVCTVTSSFSRSVSTSVGCGKN
HFSMSGSASGSFAKKTGKGGKLICKTGRYSSSKSVSAGVGKNHYSKSTSVSGSYLVAQGK
GGACTVISSFSKSVSASAGCGKNHWSKSGSCSGSYALKRGKGGKLVSETGRFSNSMSVSA
GCGKNHFSKSTSASGSYAIAQGKNGVCTVTASFSQSTSASAGCGKNHWSKSASASGSYGV
KRGKGGKLLSATGRYSSSMSASAGCGKNHWSNSASASGSYAVSLGAGQPCDA*
